# Impaired erythroid maturation in murine embryos upon loss of the preeclampsia-associated serine protease prostasin

**DOI:** 10.1101/2024.10.25.620248

**Authors:** Sara Di Carlo, Adrian Salas-Bastos, Mariela Castelblanco Castelblanco, Muriel Auberson, Marie Rumpler, Malaury Tournier, Lukas Sommer, Olaia Naveiras, Edith Hummler

## Abstract

In humans, the membrane-bound serine protease prostasin encoded by *Prss8* is associated with preeclampsia, a gestational hypertension disorder affecting blood supply of the placenta. Mice deficient in *Prss8* resulted in the death of embryos at embryonic day (E) 14.5 and it was characterized by impaired placental labyrinth maturation and vascularization. A pale phenotype was observed in these embryos, suggesting ineffective erythropoiesis. Thus, in this study we analyzed this phenotype further in *Prss8^-/-^* embryos at E11.5 and E12.5. We found a reduced number of fetal erythroblasts in placenta, yolk sac and fetal liver of *Prss8^-/-^* embryos, while the reticulocyte number was increased, suggesting a defective terminal erythroid differentiation. Further, single-cell RNA sequencing (scRNA-seq) analyses of aorta-gonad-mesonephros (AGM) revealed an upregulation of several ribosomal genes associated with Diamond-Blackfan anemia in erythroid cells of *Prss8^-/-^* (KO) embryos. These cells showed a lower capacity to maturate into erythrocytes *in vivo* and in *vitro,* despite hematopoietic cells (HSCs) being produced normally. We suggested prostasin influenced erythropoiesis in a cell-extrinsic manner, since *Prss8* expression was not detected in erythroid cells but highly expressed in ectoderm-like cells within the AGM. Congruently, while yolk sac-derived cells displayed no erythroid maturation defect *in vitro*, the yolk sac vascular remodeling in KO embryos was impaired as evidenced by reduced secondary branching likely as a consequence of the reduced blood flow. Our findings unveiled a novel role for this serine protease in terminal maturation of erythrocytes in the fetal liver and open new research avenues for understanding the physiological mechanism of prostasin and its pathological implications.

**Key Points:**

- *Prss8* deficiency causes transcriptional changes in erythroid progenitor cells in the AGM leading to impaired embryonic erythropoiesis
- Overexpression of Rpl and Rps genes by erythroid cells lacking *Prss8* leads to defective erythropoiesis and embryonic lethality

## Introduction

Proteases, and especially serine proteases, have a pivotal role in signaling processes and are thus fundamental to survival across all living organisms ^1^. Serine proteases represent the largest family of proteolytic enzymes and control several physiological processes including epithelial development, digestion, blood coagulation, inflammation and immunity. Likewise, there is growing evidence of their contribution to the pathogenesis of inflammatory and neoplastic diseases ^2,3^. Prostasin is a glycosyl-phosphatidylinositol membrane-anchored serine protease expressed in the epithelial compartment of most tissues, including prostate, lung, kidney, colon and skin, where it was shown to play a tissue-specific role ^4–6^. Prostasin was the first among many membrane-anchored serine proteases to be described as an activator of the epithelial Na^+^ channel (ENaC),^7^ and regulator of Na^+^ transport ^8,9^, thereby also named channel-activating protein 1 (CAP1). While ENaC can be proteolytically cleaved by prostasin in *in vitro* experiments,^10–12^ this has not yet been confirmed *in vivo* ^13^ Namely, prostasin has been shown to elicit a variety of non-proteolytic functions *in vivo*, such as epidermal barrier function and skin inflammation^6,14,15^,and both prostasin overexpression and downregulation have been linked to the onset of several diseases, including hypertension and cancer ^16–21^. More recently, prostasin was also found to be dysregulated in females as a sex-specific pathophysiologic difference associated to coronary microvascular dysfunction in patients with heart failure ^22^. Interestingly, *Prss8* polymorphism in human females has been linked to preeclampsia, a severe form of hypertension during pregnancy, leading to impaired placentation and premature embryonic death ^23,24^. Previous studies described a role of prostasin in human trophoblast differentiation, although its implication in embryonic development has not been yet extensively investigated ^25–28^

During embryonic development in mammals, erythropoiesis results in the rapid production of erythroid cells that support the growth and survival of the embryo and fetus ^29^. Three waves of erythropoiesis have been described to occur in the murine embryo. The first one, prior the onset of embryonic circulation at E7.5, is called “primitive” and takes place in the yolk sac giving rise to mature red blood cells (RBCs), megakaryocytes and macrophages prior the generation of hematopoietic stem cells (HSCs) ^30,31^. Concurrently, multipotential hematopoietic progenitor cells, termed “erythro-myeloid progenitors” (EMP)^32,33^ emerge in the yolk sac. These progenitors serve as major source of hematopoiesis in the developing embryo prior ^34^. *Bona fide* HSCs capable of producing all lympho-myeloid lineages arise only at late embryonic/early fetal stage of development (around E10.5) from hemogenic endothelium in the AGM. This third wave supplies adult-type erythrocytes and is thus referred to as “definitive” erythropoiesis ^29^. While primitive erythrocytes enter circulation in their nucleated form when the heartbeat starts and enucleate between E12.5 and E16.5, definitive erythrocytes circulate exclusively as enucleated cells and mature extravascularly in the fetal liver or subsequently in the bone marrow ^35^. However, the complex nature of these processes has only been elucidated in the past decades, and the precise origin of the different erythropoietic waves remains a debated field ^36–38^. In this study, we aimed to explore the physiological role of prostasin in murine embryos prior to their premature death by analyzing the phenotype and transcriptome of *Prss8^+/+^* (WT) and *Prss8^-/-^* (KO) embryos prior to E14.5. Due to the previously observed defect in vascularization and suspected impaired hemoglobinization we focused on the main hematopoietic tissues including placenta, yolk sac, AGM and fetal liver. Our data demonstrate that lack of prostasin leads to an inefficient terminal erythropoiesis likely due to an upregulation of ribonuclueoprotein (RNP)-related genes by erythroid cells found in the AGM at E11.5. This leads to impaired vessel remodeling in the yolk sac at E12.5 and embryonic lethality presumably due to anemia. Finally, our findings unwrap new research perspectives on the physiological role of prostasin as well as on the regulation of embryonic erythropoiesis.

## Methods

### Mice

*Prss8^-/-^* and *Prss8^+/+^* embryos were generated by intercrossing *Prss8**^+/-^*** mice as described previously ^39^. Mating was started in the evening, and females were checked for vaginal plugs the next morning, with the morning of plug defined as E0.5. Pregnant mice were sacrificed between 10 to 12 days later and E10.5, E11.5 and E12.5 embryos were collected for the analysis of yolk sac, AGM, fetal liver or placenta. The viability of embryos was assessed by heart beating. RNA expression of *Prss8* was performed by PCR genotyping using the following primers (*Prss8* sense: 5’- GCAGTTGTAAGCTGTCATGTG-3’; antisense (ex): 5’- TCCAGGAAGCATAGGTAGAAG-3’, sense (7): 5’- CAGCAGCTCGAGGTACCACTC -3’.

### Histology

Following dissection, yolk sac and placenta were fixed overnight in 4% PFA, hydrated in 70% ethanol and embedded in paraffin. 4 µm paraffin sections were stained with H&E to assess the general morphology. The staining was performed on the Sakura Tissue-Tek Prisma® Plus Automated Slide Stainer linked to the Tissue-Tek® Glas™ g2 Automated Glass Cover slipper. Images were acquired using the Zeiss Axioscan 7 microscope with a resolution of 40X.

### Immunofluorescence

The yolk sac was placed in 4% PFA overnight. After 24 hours, the tissue was washed 3x with PBS and blocked at RT in staining buffer (PBS + 0.3% Triton-X + 3% BSA) for 1h. It was then stained with anti-mouse CD31 (553370, BD Bioscience), anti-rat Ter119 (14-5921-82, Invitrogen) and anti-mouse VE-Cadherin (AF1002, R&D systems) antibodies overnight, followed by incubation with secondary antibodies for 1h at RT. Finally, the yolk sac was placed in mounting medium containing 4% DAPI on a glass slide and flat opened for microscopic analysis. Whole mount images were taken using a Nikon Ti2 spinning disk microscope with a resolution of 40X.

### Single cell suspension

Cells from yolk sac, AGM or fetal liver were isolated by placing each organ in one 24-well with 300 µL of dissociation medium (1mg/ml dispase-collagenase in PBS + 5mM Ca^2+^) at 37°C for 40-45 minutes. The tissues were gently crushed every 10 min using a 1000 µL pipette until complete tissue disruption. The cell suspension was collected and passed through a 40 µm cell strainer (Greiner bio-One). The collected cells were centrifuged at 1200 rpm for 5 minutes and resuspended in PBS + 10% FCS.

### Colony Forming Unit (CFU) assay

Single cell suspension was obtained as described previously and cells were counted and diluted to a final concentration of 5 x 10^5^ cells/mL. Then, 250 µL of cell suspension was added to 2.5 mL of MethoCult M34343 or M3436 medium (Stem Cell Technologies) and 1.1 mL/well was plated in duplicate in a 6 well plate. Water was added into empty spaces to minimize the medium evaporation. The read out was performed after 8 days of culture with the StemVision automated microscope (Stem Cell Technologies).

### Statistical analysis

The data shown are mean ± SEM of at least three biological replicates. Comparison between two or more groups was performed by two-tailed *t* test and values with *P* < 0.05 were considered to be statistically significant. All representations and statistical analysis were done using GraphPad Prism (version 9.3.1). Bulk RNA sequencing experiment was performed using two independent biological replicates, while single-cell RNA sequencing was performed once for each genotype and time point by pooling 3-4 embryos. Images were selected from matched embryos within a single experimental replicate to be representative of the most abundant expression pattern for each condition.

**FACS**, **imaging flow cytometry**, **bulk** and **single-cell RNA sequencing experiments** were performed, and data analyzed as described in the supplementary information.

## Results

### Vessel remodeling is impaired in the yolk sack of *Prss8^-/-^* embryos

Previous results reported embryonic lethality of *Prss8-*deficient embryos by E14.5 ^24^. To further examine the role of prostasin during embryonic development, yolk sacs from WT and KO embryos were analyzed at E12.5, just prior to premature death. The gross morphology of the yolk sac lacking prostasin showed a pale vasculature, without an evident vascular network (Figure 1A). Histological examination and CD31 immune staining of whole mount yolk sacs revealed an impaired vascular network with enlarged vessels lacking secondary branching in KO embryos (Figure 1B; Figure 2A). This difference in the vasculature was not evident at the earlier time points E10.5 and E11.5 (supplementary Figure 1A-C), indicating an essential role for prostasin during the developmental window between E11.5 and E12.5. Despite that, *Prss8* gene expression in the yolk sac of WT embryos was comparable between all-time points (supplementary Figure 1D). Next, to analyze the transcriptome of the yolk sac we performed bulk RNA sequencing (RNA-seq) at E10.5 and E11.5. Differentially expressed genes (DEGs) analysis comparing the two time points for each genotype identified a set of genes significantly and uniquely downregulated at E11.5 in the yolk sac of KO embryos (Figure 1D). Some of the downregulated genes were prevalently associated with biological processes such as angiogenesis and vasculogenesis (Figure 1E), supporting the immature vasculature observed in the phenotype of KO embryos at E12.5 (Figure 1A-B). Genes represented in the GO term vasculogenesis and angiogenesis included transcription factors and regulatory proteins, including *Tie1*, *Tek*, *Angpt1* and *Cdh5* (Figure 1F-G). However, the number of endothelial cells (CD45^-^CD43^-^; CD31^+^Ve-cad^+^) detected in the yolk sac was comparable between the two genotypes (Figure 1C), suggesting that blood vessel formation was intact. We concluded that the observed phenotype was likely not associated with defective vasculogenesis, but rather with lack of proper vessel remodeling.

**Figure 1.**
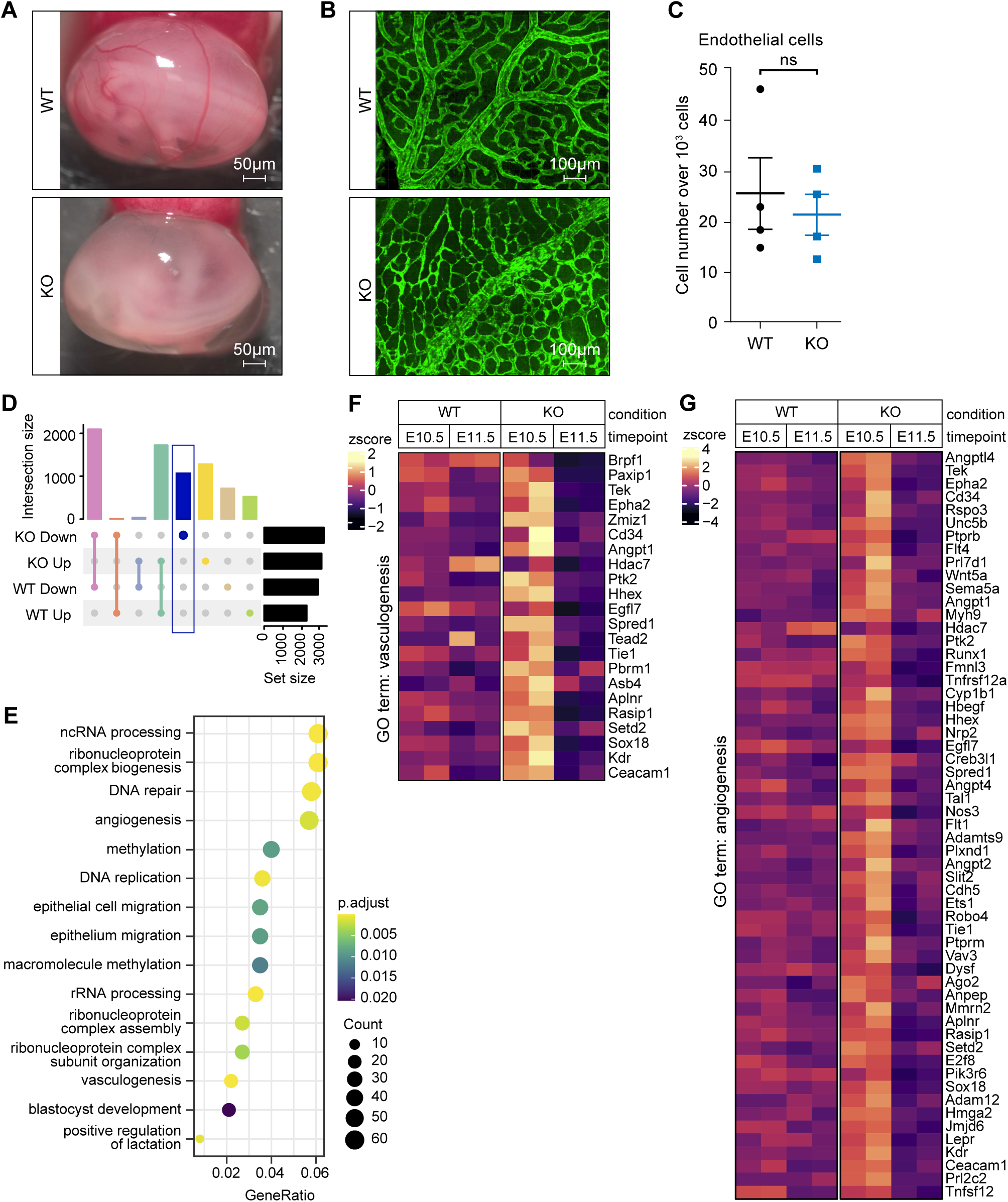
Changes in vasculogenesis-related genes correlate with impairment of vessel remodeling in the yolk sac of Prss8^-/-^ embryos. **(A**) Gross morphology of E12.5 *Prss8^+/+^* (WT) and *Prss8^-/-^* (KO) embryos; visible vessels are seen in the WT; scale bar: 50 µm. (**B**) Representative images of E12.5 whole mount yolk sac immunostaining from WT and KO embryos using antibody against CD31(GFP); scale bar: 100 µm. Magnification 40X and z-stack using a Nikon Ti2 and merged using Imaris software. (**C**) Number of endothelial cells (CD31^+^VEcad^+^) were gated on CD45^-^CD43^-^ and quantified as cell number over 10^3^ cells isolated by flow cytometry from E11.5 WT and KO yolk sac, n=4 (every dot represents one biological replicate). Fluorescent signal was recorded with the DIVA acquisition software using FACS Fortessa machine. (**D**) Upset plot of differentially down- or upregulated genes in E11.5 compared to E10.5 yolk sac from KO and WT RNA-seq data (padj < 0.05). Vertical bars show the number of shared and unique genes in each comparison. (**E)** Dot plot showing the enrichment analysis (GO biological processes) of downregulated genes in E11.5 compared to E10.5 yolk sac from KO. **(F, G**) Heatmaps showing the expression levels of DEGs related to (**F**) vasculogenesis and, (**G**) angiogenesis, respectively. Values are presented as the mean ± SEM. Significance was calculated using the unpaired Student t test. The images and staining are representative images from three biological replicates.

**Figure 2.**
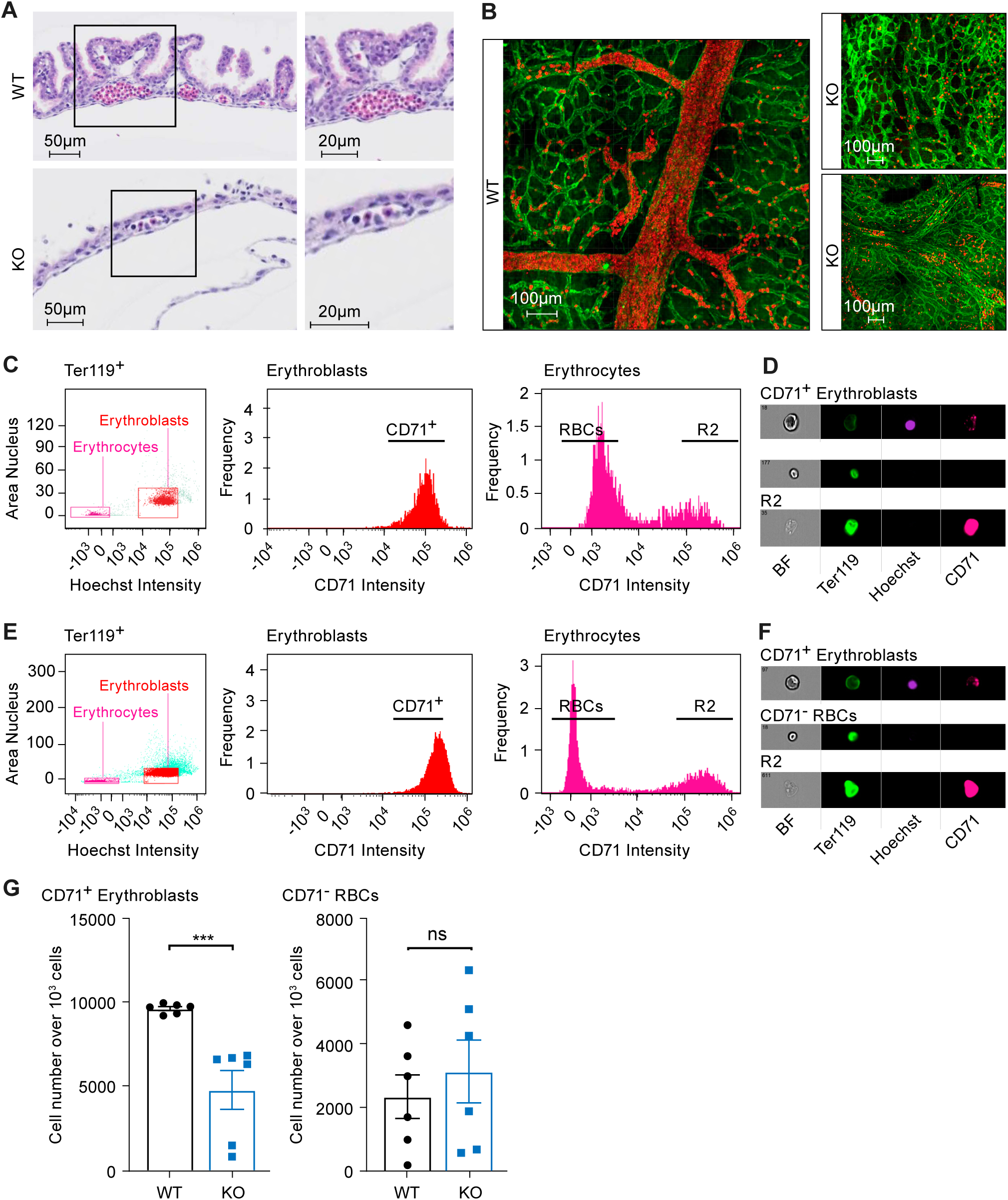
Fewer erythroblasts are found in E12.5 yolk sac from Prss8^-/-^ embryos. Representative images of **(A)** H&E-stained sections of yolk sac from WT and KO embryos; note that the visible clusters of erythrocytes (in red) are isolated, rare and present exclusively nucleated erythrocytes; scale bar: 50 µm and 20 µm, respectively. Magnification 40X, zoom using Zen Blue software, z-stack using Zeiss Axioscan 7. **(B)** Immunostaining of whole mount E12.5 yolk sac from WT and KO using antibodies against Ter119 (GFP, red) and VE-Cadherin (A647, green); scale bars: 100 µm. Magnification 40X and z-stack using a Nikon Ti2, merged using Imaris software. **(C, E)** Imaging flow cytometry analyses of yolk sac-derived cells isolated from E12.5 **(C)** WT and **(E)** KO embryos. Representative histograms of Ter119, CD71 and Hoechst-stained erythroblasts, reticulocytes and red blood cells (RBCs) are shown. **(D, F)** Images of CD71^+^ erythroblasts, CD71^-^ RBCs and dead cells (R2) cells from WT and KO embryos, respectively. Samples were acquired on ImageStreamX imaging flow cytometer (Cytek Biosciences) using DIVA acquisition software. **(G)** Quantification of indicated cell populations in **(C)** and **(E)**; n=6 biological replicates (every dot represents one biological replicate). Values are presented as the mean ± SEM. Significance was calculated using the unpaired Student t test. \*\*\**P*≤ .001. The stainings are representative images from three biological replicates.

### Defective embryonic erythropoiesis in *Prss8^-/-^* embryos

The vasculature continuously undergoes dynamic structural changes and remodeling to ensure stable vascular adaptation, which can be influenced by blood flow ^40,41^. H&E staining of the yolk sac revealed fewer nucleated erythrocytes within the vessels (Figure 2A). To further determine the cause of the vascular abnormalities, erythrocytes within the yolk sac at E12.5 were visualized by VE-cad/Ter119 immunostaining. Our results showed a profound decrease of circulating erythroid cells in yolk sacs of KO embryos (Figure 2B). This was further analyzed by imaging flow cytometry, using markers for Ter119, CD71 and Hoechst (Figure 2C and E), to distinguish the different erythroid populations. Erythroblasts exhibited a regular shape and consistent Hoechst nuclear signal, indicative of functional cells, while reticulocytes were not detected within the yolk sac of WT and KO embryos (Figure 2D and F). We found that the number of erythroblasts (Ter119^+^CD71^+^Hoechst^+^) was significantly lower in KO yolk sacs, confirming our previous observation by H&E. Whereas mature enucleated erythrocytes (Ter119^+^CD71^-^Hoechst^-^) were similar in number between the two genotypes (Figure 2G). Since the embryonic circulation has been already established at E12.5, the erythrocyte content was also analyzed in placentas at E12.5. H&E staining of placenta labyrinth from WT and KO embryos exhibited the presence of nucleated and enucleated erythrocytes (Figure 3A), as it represents the fetal-maternal interface for nutrient and gas exchange. Although placenta of KO embryos was previously described to appear pale only at E14.5^24^, quantification of the area covered by, and the number of nucleated erythrocytes showed that significantly fewer functional cells were present in KO embryos. Instead, the area covered by enucleated erythrocytes was comparable to the WT (Figure 3B). As adult erythropoiesis occurs primarily in the bone marrow leading to full enucleation, the nucleated cells are uniquely derived from embryonic circulation, indicating the embryonic red blood cells (RBCs) are fewer in the absence of prostasin, and suggesting this difference might contribute to the defective placenta development. Next, in placenta at E12.5 imaging flow cytometry using antibodies against Ter119, CD71 and Hoechst was performed to better distinguish between erythroblasts (Ter119^+^CD71^+^Hoechst^+^), mature erythrocytes (Ter119^+^Hoechst^-^CD71^-^) and reticulocytes (Ter119^+^Hoechst^-^CD71^+^) (Figure 3C and E). The cellular morphology appeared to be unaffected by the loss of prostasin (Figure 3D and F). The number of both erythroblasts and erythrocytes was significantly reduced in E12.5 KO embryos, while an accumulation of reticulocytes was observed in placentas isolated from KO embryos (Figure 3G). Taken together, these results indicate that lack of prostasin affected embryonic erythroid differentiation, beyond previously described defects in placental vasculogenesis.

**Figure 3.**
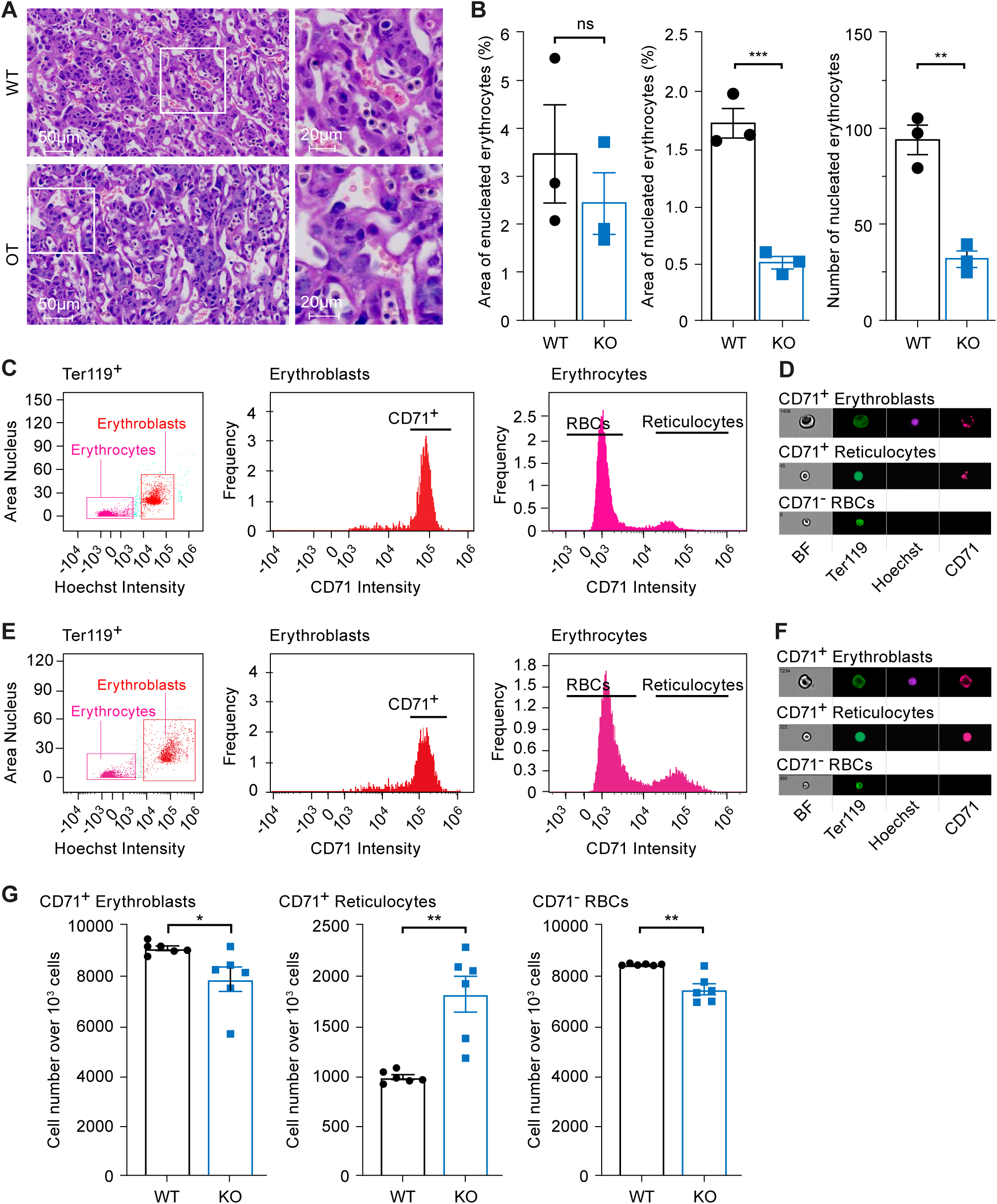
Prss8^-/-^ embryos exhibit fewer fetal erythrocytes within E12.5 placenta. **(A)** Representative H&E- stained images and their magnification of E12.5 placental labyrinths from WT and KO embryos; note the light pink enucleated erythrocytes and the dark violet nucleated ones. Scale bars: 50µm and 20µm, respectively. Magnification 40X, zoom using Zen Blue software and z-stack using Zeiss Axioscan 7. **(B)** Quantification of the area of enucleated (%, left panel) and nucleated (%, middle panel), and the number of nucleated erythrocytes (%, right panel) in placenta from WT and KO embryos. **(C, E)** Imaging flow cytometry analyses of placenta-derived cells isolated from E12.5 **(C)** WT and **(E)** KO embryos. Representative histograms of Ter119 (FITC), CD71 (Pe-Cy7) and Hoechst- stained CD71^+^ erythroblasts, CD71^+^ reticulocytes and CD71^-^ red blood cells (RBCs) are shown. **(D, F)** Images of CD71^+^ erythroblasts, reticulocytes and RBCs from WT and KO embryos, respectively. Samples were acquired on ImageStreamX imaging flow cytometer (Cytek Biosciences) using DIVA acquisition software. **(G)** Quantification of indicated cell populations in **(C)** and **(E)**; *n*= 3-6 biological replicates (every dot represents one biological replicate). Values are presented as the mean ± SEM. Significance was calculated using the unpaired Student t test. \**P*≤ .05, \*\**P* ≤ .01, \*\*\**P*≤ .001. The stainings are representative images from three biological replicates.

### Prostasin regulates terminal erythroid differentiation

Considering that developing embryonic HSCs migrate to fetal liver for clonal expansion and differentiation, imaging flow cytometry using markers for Ter119, CD71 and Hoechst was also performed in E12.5 fetal livers to distinguish and analyze the different erythroblast progenitors. The presence of maturating basophilic (BasoE), polychromatophilic (PolyE) and orthochromic (OrthoE) erythroblasts could be identified in both WT and KO embryos, while mature RBCs were absent in WT and KO embryos (Figure 4A-D). We detected a significantly reduced number of BasoE and PolyE, and in particular of OrthoE in the fetal liver of KO embryos (Figure 4A-E). Furthermore, when we analyzed the amount of lymphoid and myeloid cells present in the fetal livers, we found higher numbers of T cells, dendritic cells (DCs) and neutrophils in knockout embryos (Figure 4F; supplementary Figure 2), even though this difference was not statistically significant. This data suggests that *Prss8* deficient embryos exhibit a defective terminal erythroid differentiation, without affecting the production of other cell lineages. Next, we tested whether hematopoietic progenitor proliferation was impaired in absence of prostasin. Indeed, cells isolated from E12.5 fetal liver had lower proliferative ability *in vitro* to give rise to both erythroid (Figure 4G) and myeloid colonies (Figure 4H) in colony forming unit (CFU) assay. From the myeloid cultures we identified and quantified the largest colonies of multi-potent progenitors as colony-forming unit-granulocyte, erythroid, macrophage, megakaryocyte (CFU-GEMM). This showed that the number of GEMM colonies was unchanged between the two genotypes (Figure 4H), further suggesting that generation of multilineage progenitor cells is likely not affected. In order to investigate whether the defect in KO embryos occurred during erythroid specification or HSCs generation, the amount of hemogenic endothelium and HSC precursors was analyzed in the AGM at E11.5 by flow cytometry. The number of hemogenic endothelial cells (CD45CD43^-^; CD31Ve-cad^+^) and HSC precursors, both Type-I (CD31^+^CD41^low^CD45^-^ CD43^+^cKit^+^) and Type-II (CD31^+^CD41^low^CD45^+^CD43^+^cKit^+^), were comparable to the WT (supplementary Figure 3A-D), suggesting that HSCs were produced normally in absence of prostasin. Overall, these results indicate that *Prss8* deficiency caused an impaired specification of erythrocytes, although mRNA levels of transcription factors (TFs) that are essential for embryonic erythropoiesis in the AGM, such as *Gata1*, *Runx1* and *Tal1*, and the glycoproteins *Epo*, *Epor* were not affected (supplementary Figure 3E).

**Figure 4.**
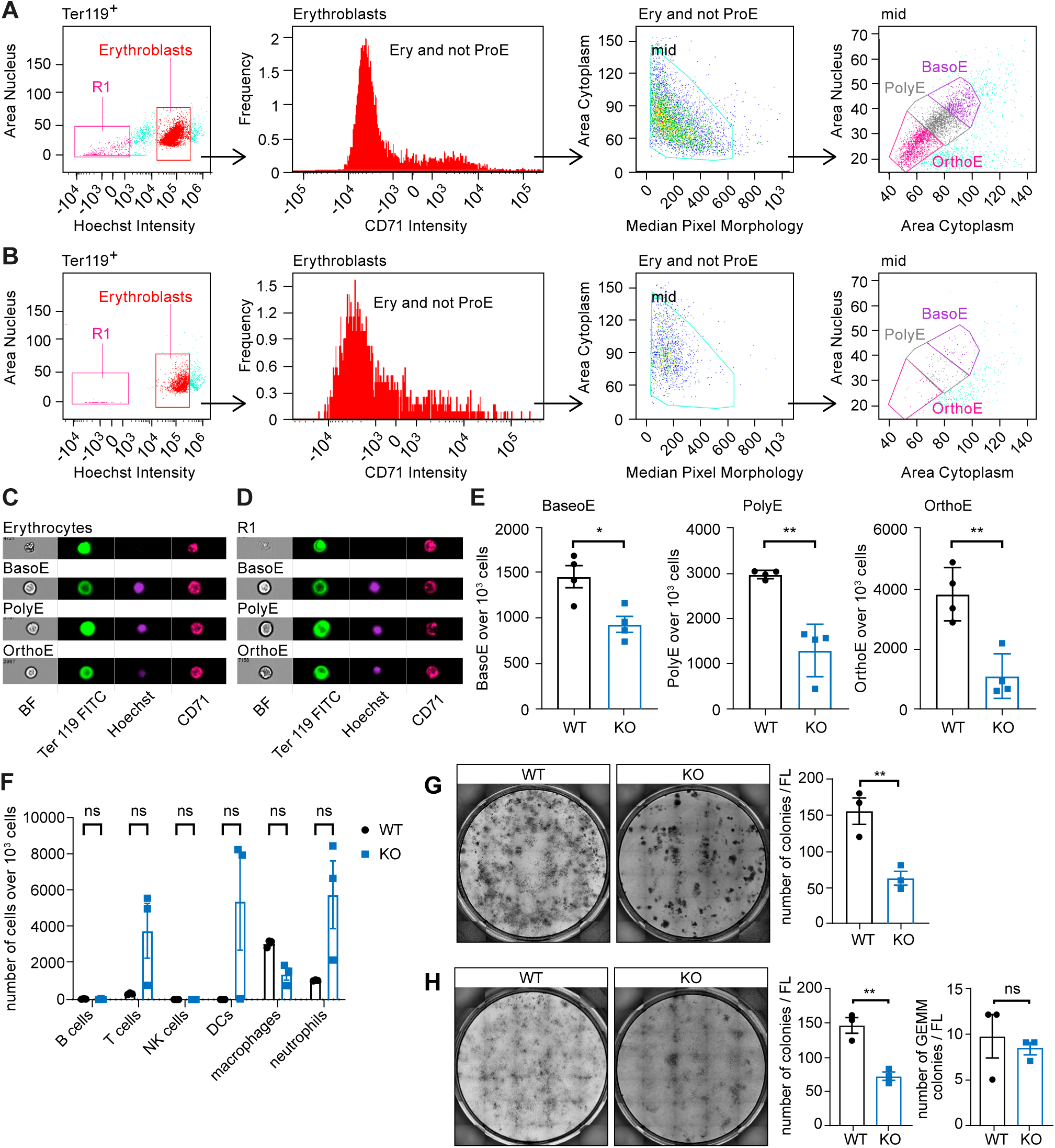
Fetal liver of E12.5 Prss8^-/-^ embryos shows a reduced number of erythroblasts. **(A, B)** Flow cytometry analyses of fetal liver-derived cells isolated from E12.5 **(A)** WT and **(B)** KO embryos. Representative histograms and gating strategy of Ter119 (FITC), CD71 (Pe-Cy7) and Hoechst-stained erythroblasts are shown. **(C, D)** Cell images of basophilic (BasoE), polychromatic (PolyE) and orthochromatic (OrthoE) erythroblasts from **(C)** WT and **(D)** KO embryos, respectively. Samples were acquired on ImageStreamX imaging flow cytometer (Cytek Biosciences) using DIVA acquisition software. **(E)** Quantification of indicated cell populations in **(A)** and **(B)**. **(F)** Number of B -, T – NK -, DC cell population as well as macrophages and neutrophils indicated as number of cells over 10^3^ fetal liver cells from WT and KO embryo. Fluorescent signal was recorded with the DIVA acquisition software using FACS Fortessa machine. **(G, H)** Colony formation unit (CFU) assays and their quantifications of fetal liver-derived cells isolated from E11.5 WT and KO embryos using **(G)** selective erythroid progenitor (MethoCult, M3436) or **(H)** hematopoietic progenitor selective medium (MethoCult, M3434); n= 3 biological replicates (every dot represents one biological replicate). Values are presented as the mean ± SEM. Significance was calculated using the unpaired Student t test. \**P*≤ .05, \*\**P* ≤ .01. The images are representative of three biological replicates.

### Erythroid cells within the AGM showed an impaired differentiation potential in absence of prostasin

To further analyze whether the impaired erythroid differentiation occurred independently of HSCs generation, we assessed the ability of AGM-derived cells at E11.5 to form erythroid or myeloid colonies using a CFU assay. In comparison to WT cells, the number of colonies derived from KO AGMs was significantly reduced in both media, indicating that overall hematopoietic progenitor activity was impaired in absence of prostasin (Figure 5A-B). Moreover, the number of GEMM-like colonies was comparable between the two genotypes, confirming that the generation of progenitor cells was not impaired by the lack of prostasin (Figure 5C). Nevertheless, despite the reduced number of erythroblasts, E11.5 yolk sac-derived cells had the colony forming capacity *in vitro* (supplementary Figure 4A-B), confirming that prostasin may play an effect in definitive erythropoiesis. The AGM is a highly interactive environment where different cell types influence erythropoiesis^42,43^. Thus, to better understand the granularity of this tissue in WT and KO embryos, we performed RNA-seq and scRNA-seq of AGM at E10.5 and E11.5. DEGs analysis of RNA-seq data comparing E10.5 and E11.5 embryos identified a set of genes significantly and uniquely downregulated in the AGM of KO embryos at E11.5 (supplementary Figure 5A). Interestingly, most of the downregulated genes were associated with microtubule organization (supplementary Figure 5B)^39^. More strikingly, entropy analysis of scRNA-seq data revealed that among the 17 clusters identified in the AGM (Figure 5D; supplementary Figure 5C-D), erythroid cluster 5 from KO embryos showed a shift toward higher entropy at E11.5 (Figure 5E-F). This might indicate an alteration in the capacity of these cells to mature between E10.5 and E11.5. Overall, these data highlight that in absence of prostasin, erythroid cells within the AGM have an impaired capacity to differentiate into erythrocytes *in vivo* and *in vitro*.

**Figure 5.**
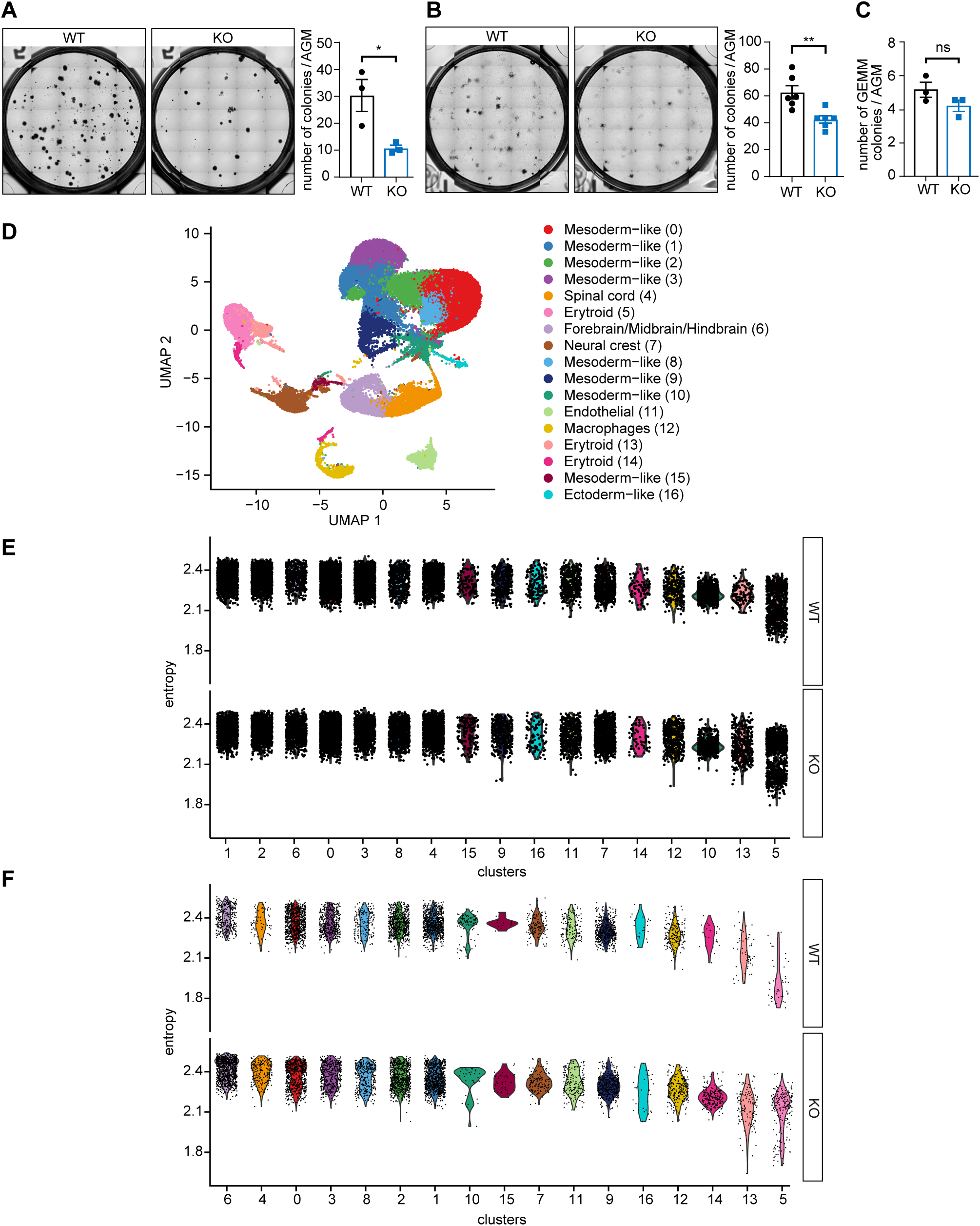
Lack of prostasin affects erythrocyte differentiation in the E11.5 AGM. **(A-C)** Colony formation unit (CFU) assays and quantifications of AGM cells isolated from E11.5 WT and KO embryos using **(A)** selective erythroid progenitor (MethoCult, M3436) or **(B, C)** hematopoietic progenitor selective medium (MethoCult, M3434), n=3 biological replicates (every dot represents one biological replicate). **(D)** Joint UMAP visualization showing annotation of all cell clusters identified by pooled scRNA-seq of AGM from E10.5 and E11.5 WT and KO embryos. **(E, F)** Analysis of cellular entropy of AGMs from E10.5 and E11.5; Values are presented as the mean ± SEM. Significance was calculated using the unpaired Student t test. \*\**P* ≤ .01. The images are representative images from three biological replicates.

### Upregulation of ribosomal protein genes linked to erythrocytes differentiation and anemia

In order to better understand the effect of *Prss8* depletion on the distinct cell populations found in the AGM, we performed DEGs analysis of scRNA-seq data. Our results revealed that the major change in gene expression occurred in the erythroid cluster 5 at E11.5, with 257 differentially expressed genes. The main upregulated pathways retrieved by the GO term analysis in E11.5 KO embryos within erythroid cluster 5 were linked to ribonucleoprotein regulation, known to influence and modulate erythropoiesis^44,45^, and biological processes including erythrocyte homeostasis and differentiation (Figure 6B). Moreover, further analysis of genes found in these pathways identified the presence of upregulated ribosomal protein small subunit (Rps) and large subunit (Rpl) genes, some of which are found to be mutated in human patients with Diamond-Blackfan anemia^46^ (Figure 6C-D; supplementary Figure 6A). This finding highlights that lack of prostasin caused changes in the transcriptome of erythroid cells, leading to their impaired differentiation already in E11.5 embryos. Interestingly, scRNA-seq analysis revealed that *Prss8* was found to be expressed only in ectoderm-like cells, while the other cell clusters including erythroid population showed no expression (supplementary Figure 6B). Moreover, in the AGM of WT embryos *Prss8* expression did not change from E10.5 to E12.5, suggesting a cell-extrinsic role of prostasin on erythropoiesis (supplementary Figure 6C). In this direction, further analysis of RNA-seq data equally indicated a significant upregulation of the GO Term “response to hypoxia” at E11.5 in both AGM and yolk sac of KO embryos, which included genes encoding critical regulators such as *Hif1a*, *Cited2*, *Ddit4* and *Pdk1* (supplementary Figure 7A-B). In conclusion, these results showed that the defective erythropoiesis observed in absence of *Prss8* is due to transcriptional changes in the erythroid cells within the AGM, which are linked to an anemic signature, likely causing embryonic lethality.

**Figure 6.**
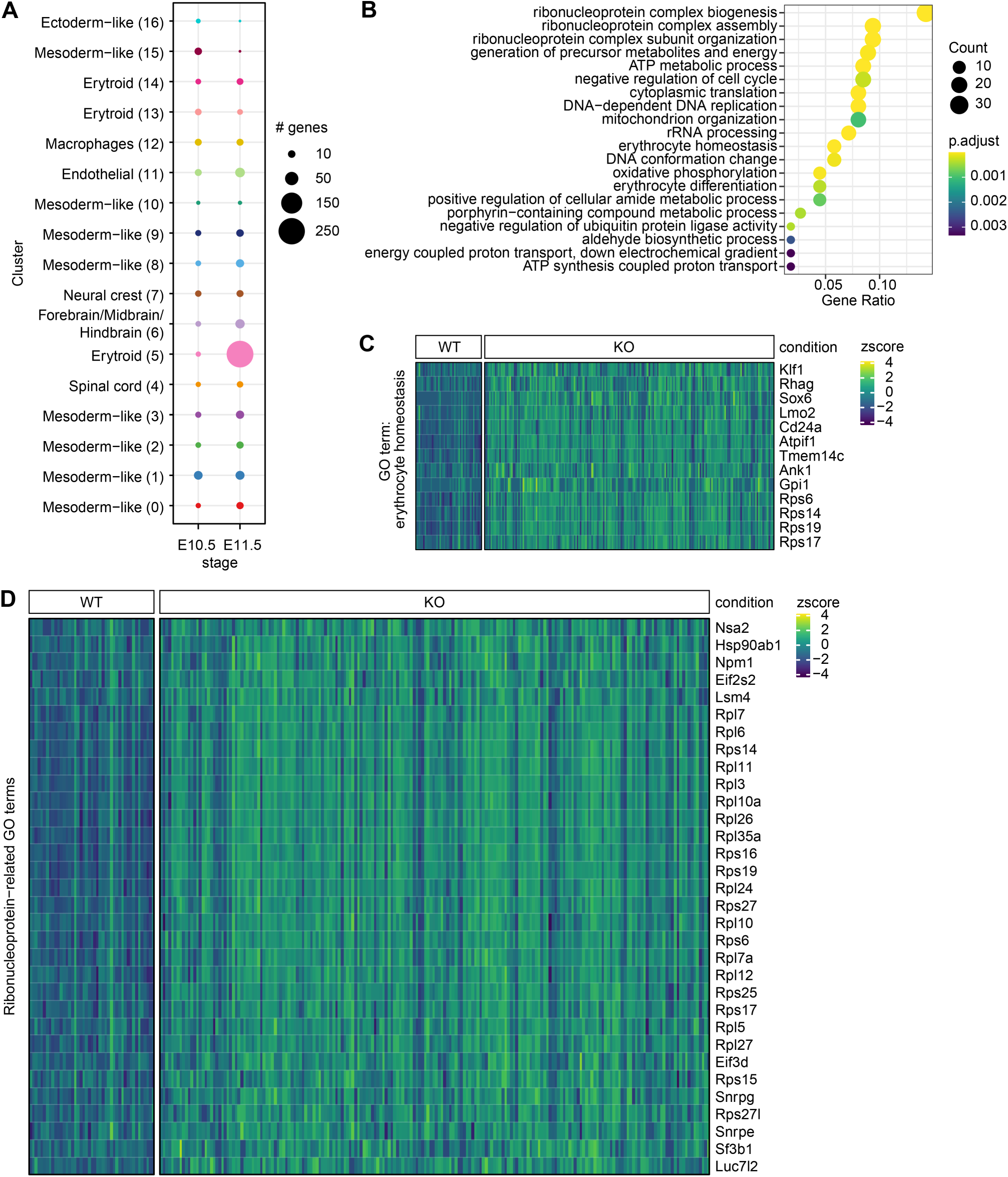
DEGs in the AGM from Prss8^-/-^ embryos are implicated in anemia and hypoxia. **(A)** Dot plot showing the number of differentially expressed genes in AGM of E10.5 and E11.5 WT and KO embryos per cluster. **(B)** GO Term analysis of upregulated genes in AGM from KO embryos at E11.5 in erythroid cluster 5. **(C, D)** Heatmaps showing the expression of **(C)** ‘erythrocyte homeostasis’ related genes (derived from B) and **(D)** ‘Ribonucleoprotein’ related genes in the erythroid cluster 5.

## Discussion

Lack of prostasin is known to cause embryonic lethality, yet its precise biological role during embryonic development remains largely unexplored ^24^. Our findings have revealed a novel function for prostasin in the regulation of erythroid differentiation in the developing embryo. We demonstrated that *Prss8* deficiency resulted in transcriptional changes of AGM-isolated erythroid cells, which explains their impaired capacity to give rise to mature erythrocytes. The defective fetal liver erythropoiesis resulted in an aberrant remodeling of the yolk sac vasculature and anemia, which likely contributes to impaired placental vascularization and lethality of *Prss8^-/-^* embryos.

We found the defective vessel remodeling of the yolk sac to be associated with a downregulation of key regulators of angiogenesis and vasculogenesis. Interestingly, some of the downregulated genes namely *Brpf1, Angpt1, Hdac7 and Setd2*, have already been reported to cause embryonic lethality if deleted, showing a similar phenotype as *Prss8^-/-^* embryos ^47–50^. Although these proteins were described to regulate vessel development, our data suggest that the vascular phenotype observed in *Prss8^-/-^* embryos was not associated with defective vasculogenesis, since blood vessel formation was intact. Instead, we found differences in the number of erythroblasts at E12.5, which were significantly reduced in placenta, yolk sac and fetal liver of *Prss8^-/-^*embryos. We reasoned that this difference is tissue independent as a result of free cells circulating between extra- and intra- embryonic compartments from E10 onwards ^51^. Erythroblasts are known to further differentiate extravascularly in fetal liver ^52^, yet reticulocytes were absent in the fetal liver of E12.5 WT and KO embryos (data not shown). Instead, we observed an accumulation of reticulocytes and a reduction of mature erythrocytes in placenta of E12.5 *Prss8^-/-^* embryos, suggesting that the placenta is a crucial site of erythrocyte maturation even as the fetal liver becomes a hematopoietic organ. These findings are in line with other studies showing that erythrocyte content and blood flow influence vessel remodeling ^41,53,54^.

Starting from E10.5 erythroblasts arise from HSC precursors, which are common progenitors for all blood cells ^55^. We showed that HSCs generation in the AGM was not affected by the loss of prostasin, despite the reduced number of erythroblasts. This finding was confirmed by FACS analysis of the fetal liver, where we observed a slightly increased number of T cells, DCs and neutrophils in knockout embryos, which is in line with studies in human patients with severe anemia^56–59^. Further, we observed that the number of erythroid colonies from AGM and fetal liver culture *in vitro* was reduced in *Prss8^-/-^* embryos, but not the number of GEMM- like progenitors, further indicating that the implication of prostasin in the differentiation of other cell lineages was unlikely. Overall, the functionality of HSCs was maintained. Although the number of erythroid and myeloid colonies was significantly reduced in KO cells, we speculate that this can be a result of erythroid colonies not appearing. Yet no differences were observed in yolk sac culture, suggesting that prostasin acts specifically on the maturation of the definitive erythroid lineage in the fetal liver. However, we could not fully exclude the contribution of mobilized progenitors.

Identification of the erythropoietic niche and characterization of its environment has been the focus of many research studies over the past years. Sampling by scRNA-seq allows monitoring of these processes with minimal manipulation of the milieu^60,61^. We were able to show that in the AGM, the major change in gene expression between *Prss8^+/+^* and *Prss8^-/-^*embryos at E11.5 occurred in the erythroid cells, and this might explain the poorly differentiated erythrocytes in *Prss8^-/-^* embryos. Interestingly, in the dataset many of the upregulated genes in KO embryos were related to ribonucleoprotein complex regulation. This result is in line with other studies reporting that mutation of ribosomal protein genes is linked with defective erythroid maturation and anemia^44,45,62^. Notably, we also found several Rpl and Rps genes mutated in patients with Diamond-Blackfan anemia to be upregulated in erythroid cells of *Prss8^-/-^* embryos. This might indicate that immature erythroid cells failed to differentiate due to overexpression of Rpl and Rps genes. In the AGM of E11.5 *Prss8^-/-^* embryos, RNA-seq data revealed a downregulation of genes related to microtubule complex organization, a process known to be critical for erythrocyte maturation ^63–65^. Additionally, we found that genes related to hypoxic signature were upregulated in yolk sac and AGM of E11.5 *Prss8 ^-/-^* embryos, which may be due to lower levels of oxygen in those tissues.

In the AGM, *Prss8* was not detectable in erythroid cells but mainly in the ectoderm, suggesting that prostasin could cause changes in the transcriptome of erythroid cells in a cell-extrinsic manner. Considering ectoderm-derived cells as important constituents of the niche, which support hematopoiesis in adult bone marrow and embryonic AGM ^66,67^, we propose that it might also influence erythropoiesis in the fetal liver. Further investigations are required to confirm this hypothesis, and conditional HSCs knockouts as well as co-culture experiments might help to further unveil the role of *Prss8* during fetal liver erythropoiesis. In conclusion, while the main target of prostasin still needs to be identified, our data highlight that *Prss8* controls murine definitive erythropoiesis between E10.5 and E11.5 through erythroid-specific expression of RPs genes. This warrants further studies on prostasin and may provide novel insights for erythropoiesis-related disorders. Moreover, the developmental defect of the placental vasculature at E14.5 associated with *Prss8^-/-^* embryonic lethality and onset of preeclampsia ^23,24^ might thus occur as a consequence of reduced embryonic erythropoiesis.

## Data Sharing Statement

- All original data are available upon reasonable request from the corresponding author Sara Di Carlo
- The RNA sequencing data from this study have been deposited in the GEO data repository under the accession numbers GSE275398 (https://www.ncbi.nlm.nih.gov/geo/query/acc.cgi?acc=GSE275398) and GSE275399 (https://www.ncbi.nlm.nih.gov/geo/query/acc.cgi?acc=GSE275399).

## Acknowledgements

The authors would like to thank the members of the Hummler and Naveiras laboratories for useful discussions and technical support. We also acknowledge the excellent graphical work by Mia Braunwalder (Zürich), the Histology Core Facility of the Ecole Polytechnique Fédérale de Lausanne (EPFL) for their technical input of H&E staining, the Lausanne Genomics Technologies Facility (GTF, UNIL) from the University of Lausanne for the bulk- and single-cell RNA sequencing experiments, and the Cellular Imaging Facility (CIF, UNIL) for the microscope use and guidance.

This research has been funded by Swiss National Science Foundation (grant 31003A-182478/1 and 31003A-163347) to E.H.

The visual abstract was created using BioRender.com.

## Authorship

Contribution: E Hummler, M Castelblanco and S Di Carlo designed the work; M Castelblanco, M Auberson, M Rumpler, M Tournier and S Di Carlo performed the experiments and analyzed the data, A Salas-Bastos and L Sommer provided useful insights and performed bioinformatic analysis of the data; E Hummler, O Naveiras and S Di Carlo supervised the work and interpreted the data; S Di Carlo wrote the manuscript; E Hummler and O Naveiras provided critical comments on the manuscript.

Disclosure of Conflict of Interest: the authors declare that they have no conflict of interest.

## Figure legends

**Supplementary Figure 1.**
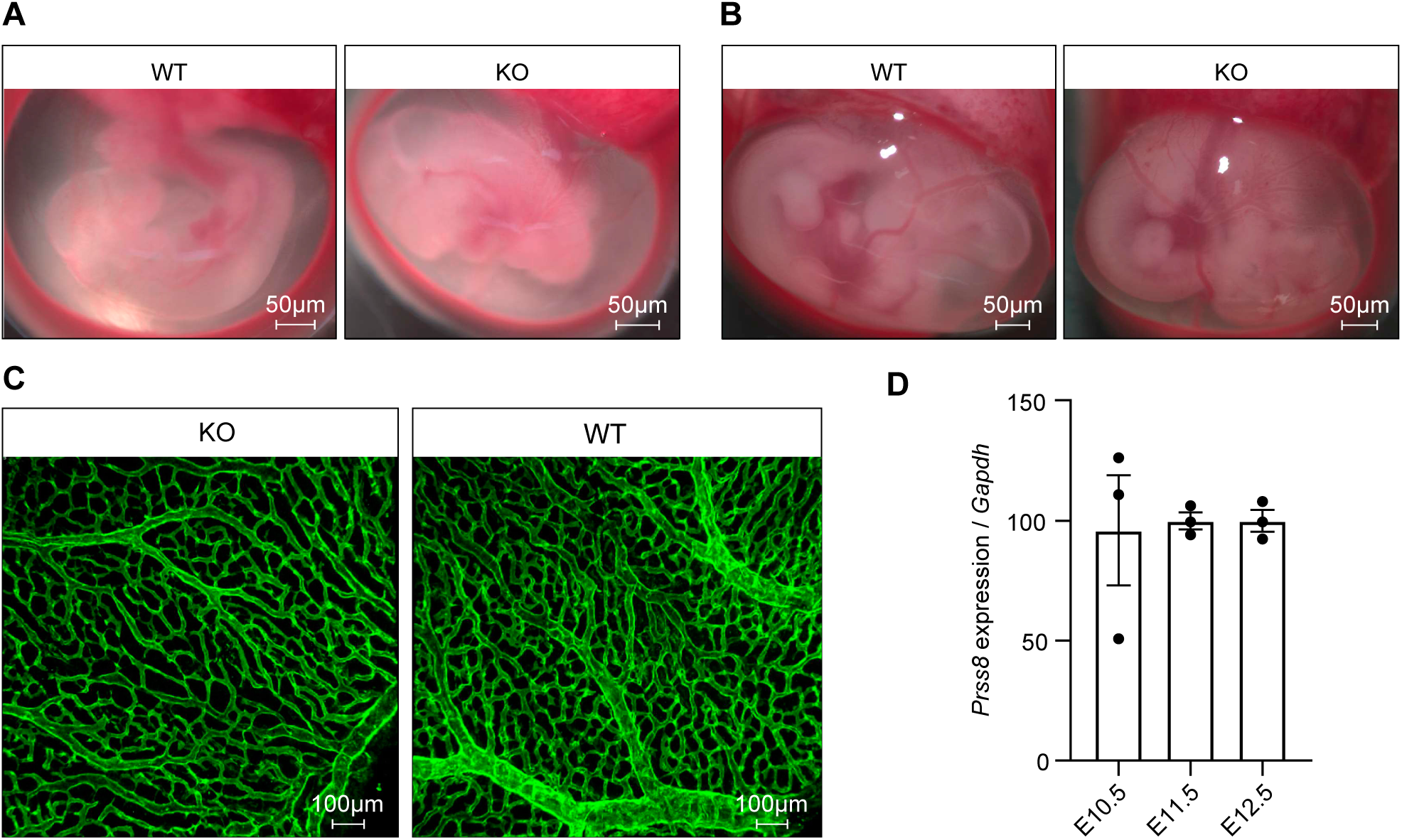
Similar phenotype and vasculature of the yolk sac of Prss8^-/-^ embryos at E10.5 and E11.5. Gross morphology of **(A)** E10.5 and **(B)** 11.5 WT and KO embryos; Scale bar: 50 µm. **(C)** CD31 immunostaining of whole mount yolk sac from E11.5 wild-type and knockout embryos; **(D)** qRT-PCR analysis of *Prss8* expression in yolk sac from WT at E10.5, E11.5 and E12.5; n=3, values are presented as the mean ± SEM. Significance was calculated using the unpaired Student t test. The images are representative images from three biological replicates.

**Supplementary Figure 2.**
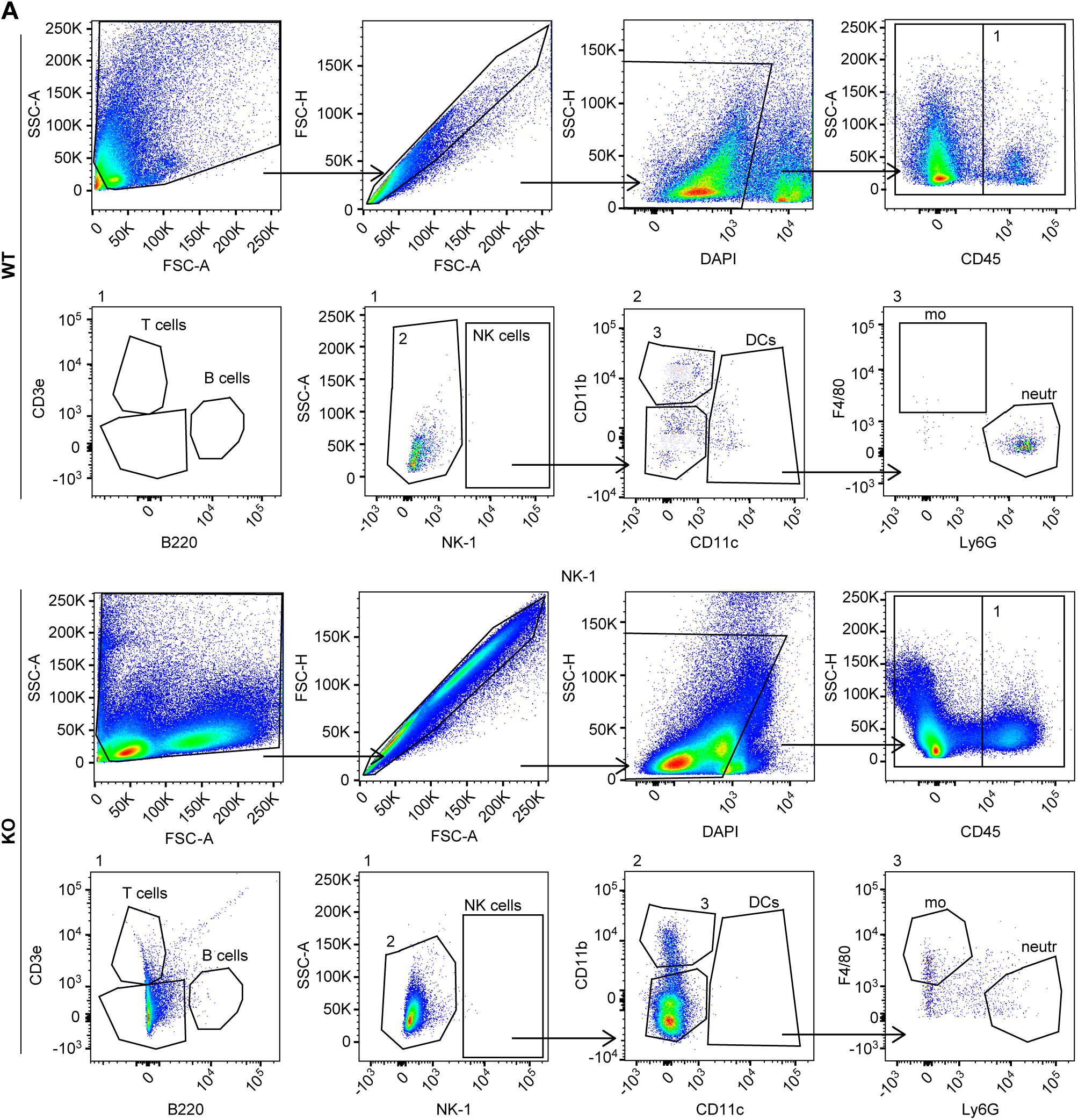
Gating strategy for myeloid and lymphoid cells in E12.5 fetal liver. **(A)** Representative plots of flow cytometry analysis using antibodies against CD45 (BUV737), B220 (BV711), NK-1 (FITC), CD11b (BV421), CD11c (PerCP/Cy5.5), F4/80 (PE-Texas red) and Ly6G (APC/Cy7) on cells isolated from WT (upper panel) and KO (bottom panel) embryos; *n*=3, the plots are representative images from three biological replicates. Fluorescent signal was recorded with the DIVA acquisition software using FACS Fortessa machine.

**Supplementary Figure 3.**
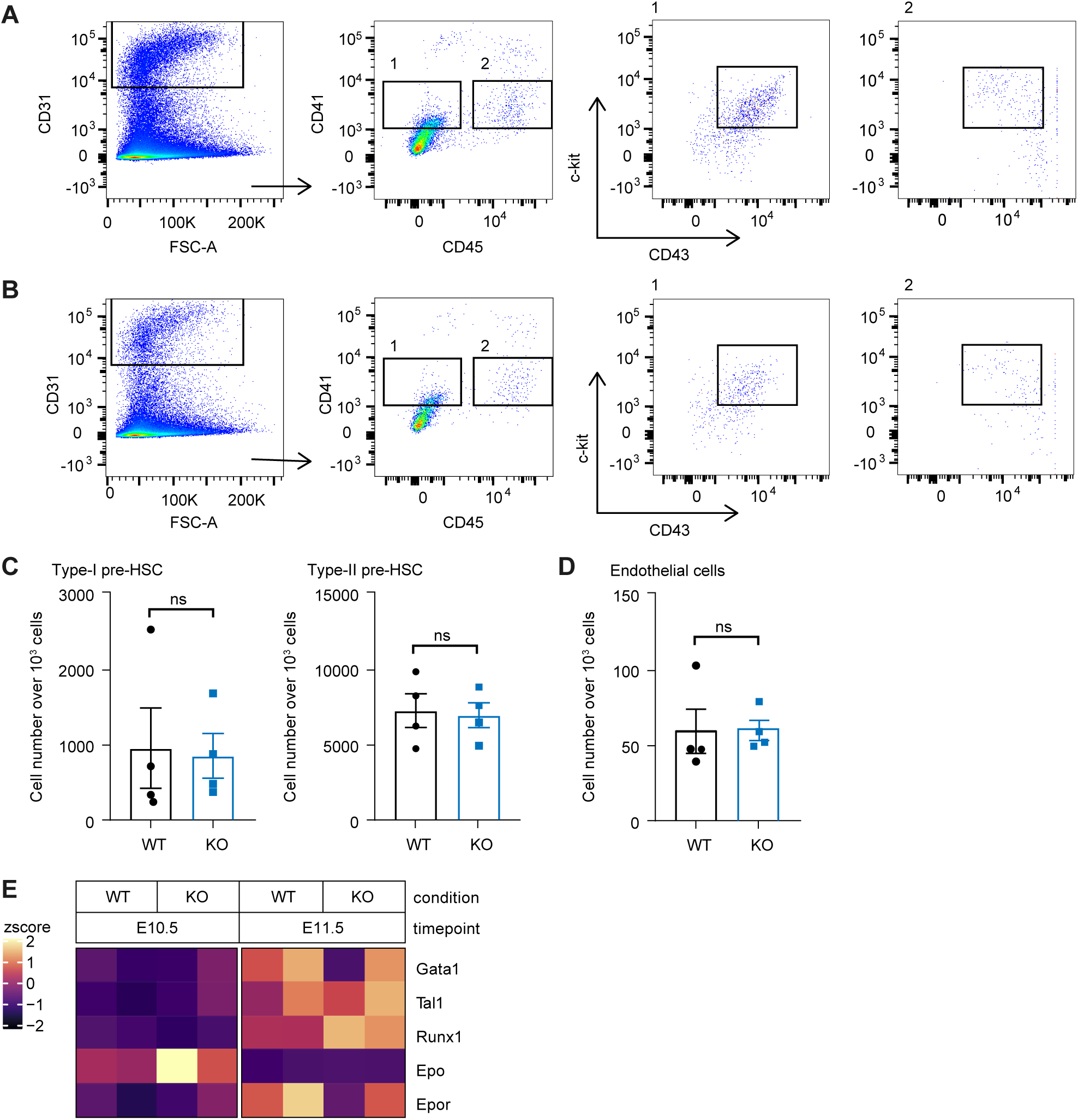
Gating strategy for HSCs and gene expression of erythropoiesis genes in E11.5 AGM. **(A-C)** Gating strategy **(A, B)** and quantification **(C)** using flow cytometry of AGM-derived cells from E11.5 **(A)** WT and **(B)** KO embryos identifying Type I and Type II pre-HSC cells using antibodies against CD45 (BUV 737), CD41 (FITC), CD43 (PE7Cy7) and CD31 (PE); n=4 (every dot represents a biological replicate). Fluorescent signal was recorded with the DIVA acquisition software using FACS Fortessa machine. (**D**) Quantification of endothelial cells expressed as cell number over 10^3^ total cells from WT and KO embryos; n=4 (every dot represents a biological replicate). **(E)** Heatmap showing the expression levels of *Gata1*, *Tal1, Runx1, Epo, Epor* in E10.5 and E11.5 WT and KO embryos, respectively. The stainings are representative images from three biological replicates. Values are presented as the mean ± SEM. Significance was calculated using the unpaired Student t test.

**Supplementary Figure 4.**
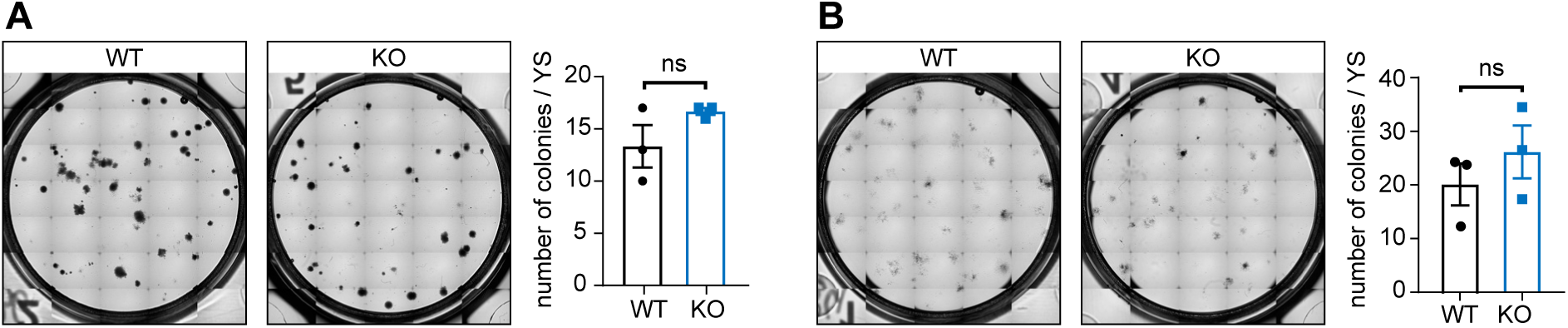
*In vitro* culture of E11.5 yolk sac-derived cells. Colony formation unit (CFU) assays and their quantifications from WT and KO embryos using **(A)** selective erythroid progenitor (MethoCult, M3436) or **(B)** hematopoietic progenitor selective medium (MethoCult, M3434). n=3, values are presented as the mean ± SEM. Significance was calculated using the unpaired Student t test. The images are representative images from three biological replicates.

**Supplementary Figure 5.**
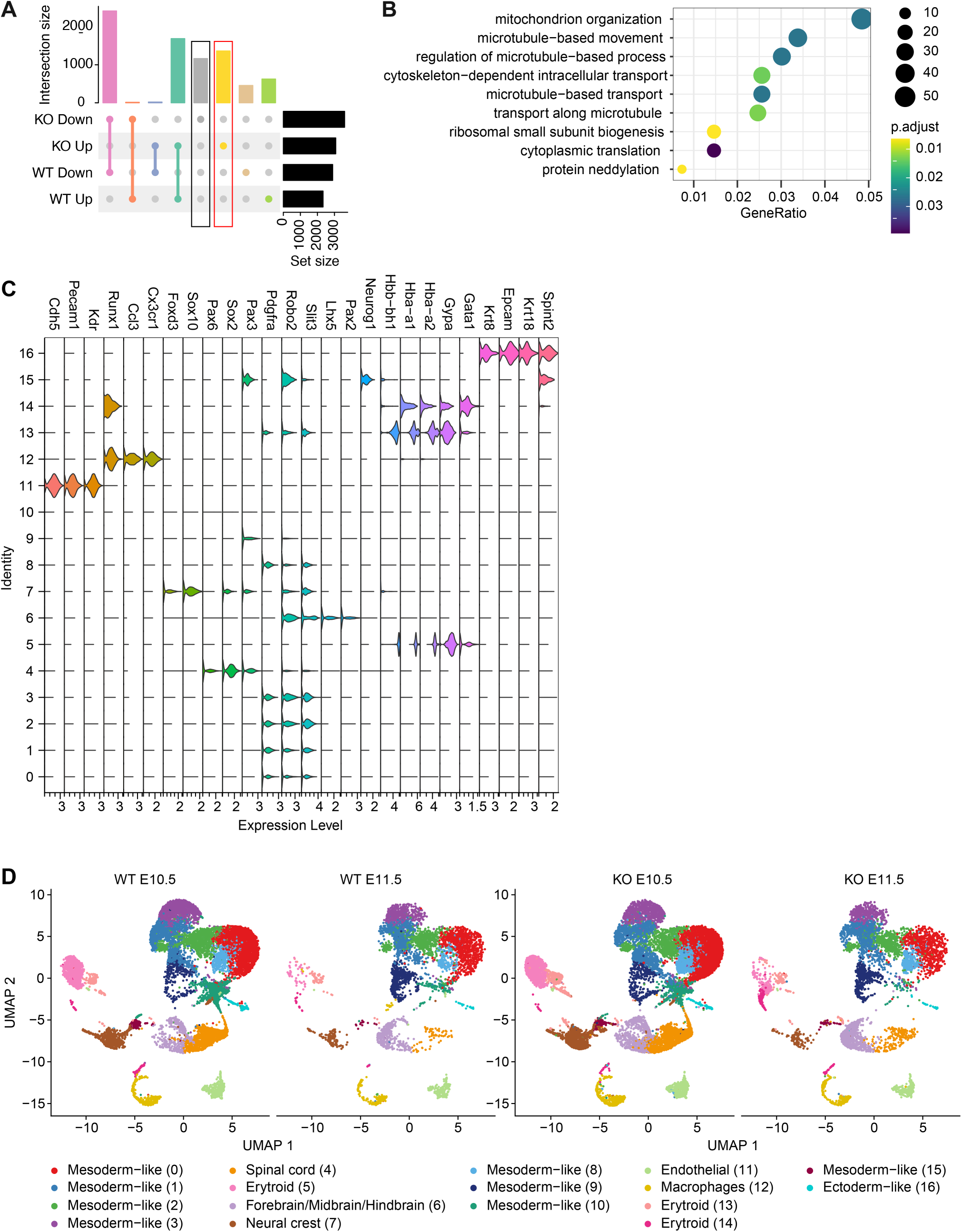
E11.5 AGM from Prss8^-/-^ embryos shows reorganization of microtubules. **(A)** Upset plot of differentially down- or upregulated genes in E11.5 versus E10.5 AGM from KO and WT (padj < 0.05). Vertical bars show the number of shared and unique genes in each comparison. **(B)** Dot plot showing the enrichment analysis (GO biological processes) of upregulated genes in E11.5 versus E10.5 AGM from KO embryos (yellow bar in A). **(C)** Violin plot of representative gene markers used to identify cells clusters in AGM samples of scRNA-seq data. **(D)** Heatmap showing the expression levels of genes related to the GO term ‘erythrocyte differentiation’ (derived from Figure 6B) in the erythroid cluster 5 of E11.5 WT and KO embryos.

**Supplementary Figure 6.**
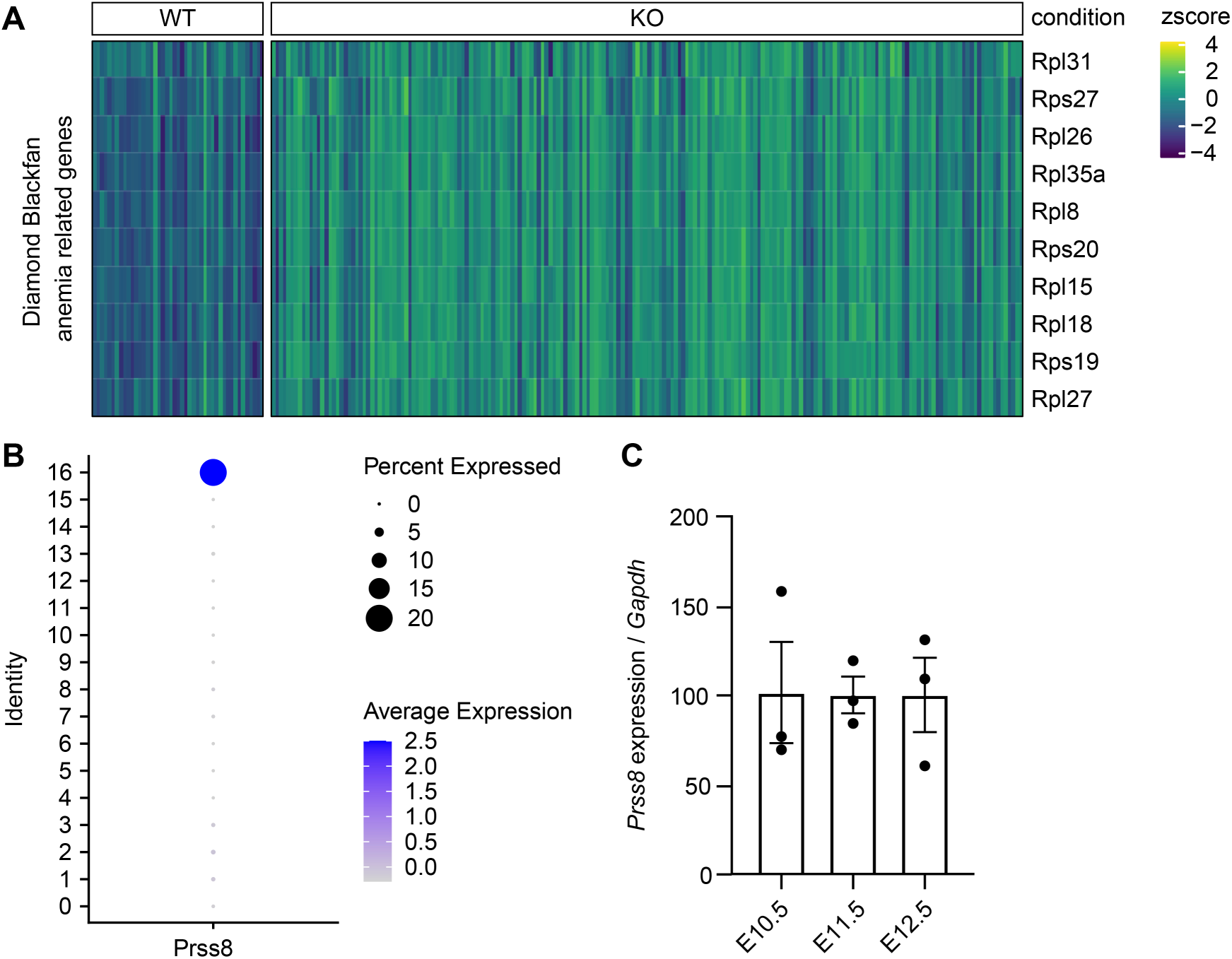
*Prss8* is expressed mainly by ectoderm-like cells in the AGM. **A)** Heatmap of significantly upregulated genes related to ‘Diamond-Blackfan’ anemia’ in KO AGMs at E11.5 from scRNA-seq analysis; **(B)** Dot plot showing Prss*8* expression in the distinct cell clusters of WT AGMs E11.5 scRNA- seq analysis. **(C)** qRT-PCR analysis of *Prss8* expression in AGM from WT embryos at E10.5, E11.5 and E12.5. n= 3, values are presented as the mean ± SEM. Significance was calculated using the unpaired Student t test.

**Supplementary Figure 7.**
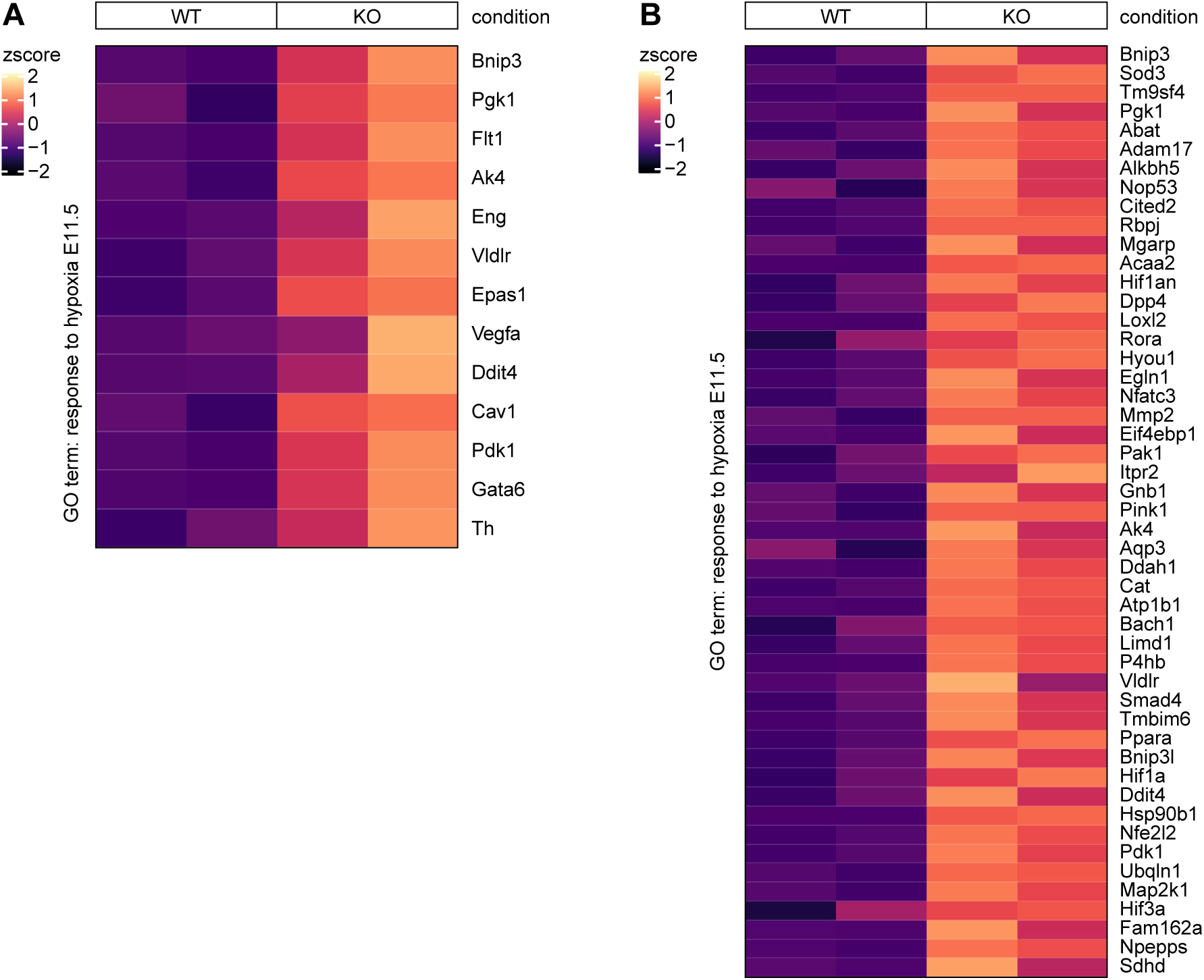
Expression of hypoxia related genes in AGM and yolk sac at E11.5. **(A, B)** Heatmaps showing the expression of ‘response to hypoxia’ related genes in E11.5 **(A)** AGM and **(B)** yolk sac of WT and KO embryos from bulk RNA-seq analysis.

## References

1. Patel S. A critical review on serine protease: Key immune manipulator and pathology mediator. Allergol Immunopathol (Madr). 2017;45(6):579–591.

2. Antalis TM, Bugge TH, Wu Q. Membrane-anchored serine proteases in health and disease. Prog Mol Biol Transl Sci. 2011;99(C):1–50.

3. Szabo R, Bugge TH. Membrane-anchored serine proteases in vertebrate cell and developmental biology. Annu Rev Cell Dev Biol. 2011;213–235.

4. Malsure S, Wang Q, Charles RP, et al. Colon-specific deletion of epithelial sodium channel causes sodium loss and aldosterone resistance. J Am Soc Nephrol. 2014;25(7):1453– 1464.

5. Planes C, Randrianarison NH, Charles RP, et al. ENaC-mediated alveolar fluid clearance and lung fluid balance depend on the channel- activating protease 1. EMBO Mol Med. 2010;2(1):26–37.

6. Frateschi S, Camerer E, Crisante G, et al. PAR2 absence completely rescues inflammation and ichthyosis caused by altered CAP1/Prss8 expression in mouse skin. Nat Commun. 2011;2–161.

7. Rossier BC, Jackson Stutts M. Activation of the epithelial sodium channel (ENaC) by serine proteases. Annu Rev Physiol. 2009;71:361–379.

8. Anand D, Hummler E, Rickman OJ. ENaC activation by proteases. Acta Physiologica. 2022;235(1):e13811.

9. Aufy M, Hussein AM, Stojanovic T, Studenik CR, Kotob MH. Proteolytic activation of the epithelial sodium channel (ENaC): its mechanisms and implications. Int. J. Mol. Sci. 2023;24:17563.

10. Bruns JB, Carattino MD, Sheng S, et al. Epithelial Na+ channels are fully activated by furin- and prostasin-dependent release of an inhibitory peptide from the γ-subunit. Journal of Biological Chemistry. 2007;282(9):6153–6160.

11. Adachi M, Kitamura K, Miyoshi T, et al. Activation of epithelial sodium channels by prostasin in xenopus oocytes. Journal of the American Society of Nephrology. 2001;12(6):1114–1121.

12. Carattino MD, Mueller GM, Palmer LG, et al. Prostasin interacts with the epithelial Na channel and facilitates cleavage of the γ subunit by a second protease. Am J Physiol Renal Physiol. 2014;307:1080–1087.

13. Ehret E, Stroh S, Auberson M, et al. Kidney-specific membrane-bound serine proteases CAP1/Prss8 and CAP3/St14 affect ENaC subunit abundances but not its activity. Cells. 2023;12(19):2342.

14. Crisante G, Battista L, Iwaszkiewicz J, et al. The CAP1/Prss8 catalytic triad is not involved in PAR2 activation and protease nexin-1 (PN-1) inhibition. The FASEB Journal. 2014;4792– 805.

15. Svenningsen P. Non-enzymatic function of prostasin and sodium balance. Acta Physiologica. 2021;232(1):e13649.

16. Takahashi S, Suzuki S, Inaguma S, et al. Down-regulated expression of prostasin in high- grade or hormone-refractory human prostate cancers. Prostate. 2003;54:187–193.

17. Chen L-M, Hodge GB, Guarda LA, et al. Down-Regulation of Prostasin Serine Protease: A Potential Invasion Suppressor in Prostate Cancer. Prostate. 2001;48:93–103.

18. Chen L-M, Chai KX. Prostasin serine protease inhibits breast cancer invasiveness and is transcriptionally regulated by promoter DNA methylation. Int. J. Cancer. 2002;97:323– 329.

19. Maekawa A, Kakizoe Y, Miyoshi T, et al. Camostat mesilate inhibits prostasin activity and reduces blood pressure and renal injury in salt-sensitive hypertension. J Hypertens. 2009;27(1):181–189.

20. Koda A, Wakida N, Toriyama K, et al. Urinary prostasin in humans: relationships among prostasin, aldosterone and epithelial sodium channel activity. Hypertension Research. 2009;276–281.

21. Olivieri O, Castagna A, Guarini P, et al. Urinary prostasin a candidate marker of epithelial sodium channel activation in humans. Hypertension. 2005;683–688.

22. Chandramouli C, Ting TW, Tromp J, et al. Sex differences in proteomic correlates of coronary microvascular dysfunction among patients with heart failure and preserved ejection fraction. Eur J Heart Fail. 2022;24:681–684.

23. Ejaz S, Ali A, Azim K, et al. Association between preeclampsia and prostasin polymorphism in pakistani females. Saudi Med J. 2020;41(11):1234–1240.

24. Hummler E, Dousse A, Rieder A, et al. The channel-activating protease CAP1/Prss8 is required for placental labyrinth maturation. PlosOne. 2013;8(2):e55796.

25. Yang Y, Zhang J, Gong Y, et al. Increased expression of prostasin contributes to early-onset severe preeclampsia through inhibiting trophoblast invasion. Journal of Perinatology. 2015;35:16–22.

26. Fu Y-Y, Gao W-L, Chen M, et al. Prostasin regulates human placental trophoblast cell proliferation via the epidermal growth factor receptor signaling pathway. Human Reproduction. 2010;25(3):623–632.

27. Szabo R, Molinolo A, List K, Bugge TH. Matriptase inhibition by hepatocyte growth factor activator inhibitor-1 is essential for placental development. Oncogene. 2007;26(11):1546–1556.

28. Frateschi S, Keppner A, Malsure S, et al. Mutations of the serine protease CAP1/Prss8 lead to reduced embryonic viability, skin defects, and decreased ENaC activity. American Journal of Pathology. 2012;181(2):605–615.

29. Baron MH, Isern J, Fraser ST. The embryonic origins of erythropoiesis in mammals. Blood. 2012;119(21):4828–4837.

30. Soares-Da-Silva F, Freyer L, Elsaid R, et al. Yolk sac, but not hematopoietic stem cell– derived progenitors, sustain erythropoiesis throughout murine embryonic life. Journal of Experimental Medicine. 2021;218(4):e20201729.

31. Kingsley PD, Malik J, Fantauzzo KA, Palis J. Yolk sac-derived primitive erythroblasts enucleate during mammalian embryogenesis. Blood. 2004;104(1):19–25.

32. Lacaud G, Gore L, Kennedy M, et al. Runx1 is essential for hematopoietic commitment at the hemangioblast stage of development in vitro. Blood. 2002;100(2):458–466.

33. Palis J, Chan RJ, Koniski A, et al. Spatial and temporal emergence of high proliferative potential hematopoietic precursors during murine embryogenesis. PNAS. 2001;98(8):4528–4533.

34. Frame JM, McGrath KE, Palis J. Erythro-myeloid progenitors: “Definitive” hematopoiesis in the conceptus prior to the emergence of hematopoietic stem cells. Blood Cells Mol Dis. 2013;51(4):220–225.

35. Ji RP, Phoon CKL, Aristizábal O, et al. Onset of cardiac function during early mouse embryogenesis coincides with entry of primitive erythroblasts into the embryo proper results UBM-doppler imaging of early-stage embryos initiation of early cardiovascular function and correlation with erythroblast entry. Circ Res. 2003;92:133–135.

36. Ganuza M, Chabot A, Tang X, et al. Murine hematopoietic stem cell activity is derived from pre-circulation embryos but not yolk sacs. Nat Commun. 2018;9(1):5405.

37. Yumine A, Fraser ST, Sugiyama D. Regulation of the embryonic erythropoietic niche: A future perspective. Blood Res. 2017;52(1):10–17.

38. Stefanska M, Batta K, Patel R, et al. Primitive erythrocytes are generated from hemogenic endothelial cells. Sci Rep. 2017;7(1):6401.

39. Rubera I, Meier E, Vuagniaux G, et al. A conditional allele at the mouse channel activating protease 1 (Prss8) gene locus. Genesis. 2002;32(2):173–176.

40. Kasaai B, Caolo V, Peacock HM, et al. Erythro-myeloid progenitors can differentiate from endothelial cells and modulate embryonic vascular remodeling. Sci Rep. 2017;7:43817.

41. Lucitti JL, Jones EAV, Huang C, et al. Vascular remodeling of the mouse yolk sac requires hemodynamic force. Development. 2007;134(18):3317–3326.

42. Rybtsov S, Sobiesiak M, Taoudi S, et al. Hierarchical organization and early hematopoietic specification of the developing HSC lineage in the AGM region. Journal of Experimental Medicine. 2011;208(6):1305–1315.

43. Todorovic V. It takes a village to raise a hematopoietic stem cell. Nature Cardiovascular Research. 2022;1(9):793.

44. Zheng Z, Yang S, Gou F, et al. The ATF4-RPS19BP1 axis modulates ribosome biogenesis to promote erythropoiesis. Blood. 2024;144(7):742–756.

45. Chennupati V, Veiga DFT, Maslowski KM, et al. Ribonuclease inhibitor 1 regulates erythropoiesis by controlling GATA1 translation. Journal of Clinical Investigation. 2018;128(4):1597–1614.

46. Member of the MaRIH rare diseases health network. French National Diagnosis and Care Protocol (PNDS): acquired and inherited aplastic anemia. 2023.

47. Fengling Chen, Jiewen Chen, Hong Wang, et al. Histone lysine methyltransferase SETD2 regulates coronary vascular development in embryonic mouse hearts. Front Cell Dev Biol. 2021;9:1–12.

48. Chang S, Young BD, Li S, et al. Histone deacetylase 7 maintains vascular integrity by repressing matrix metalloproteinase 10. Cell. 2006;126(2):321–334.

49. Jeansson M, Gawlik A, Anderson G, et al. Angiopoietin-1 is essential in mouse vasculature during development and in response to injury. Journal of Clinical Investigation. 2011;121(6):2278–2289.

50. You L, Yan K, Zou J, et al. The chromatin regulator Brpf1 regulates embryo development and cell proliferation. J Biol Chem. 2015;290(18):11349–11364.

51. Dzierzak E, Speck NA. Of lineage and legacy: the development of mammalian hematopoietic stem cells. Nat Immunol. 2008;9(2):129–136.

52. Palis J. Primitive and definitive erythropoiesis in mammals. Front Physiol. 2014;5(3):00003.

53. Kasaai B, Caolo V, Peacock HM, et al. Erythro-myeloid progenitors can differentiate from endothelial cells and modulate embryonic vascular remodeling. Sci Rep. 2017;7:43817.

54. Goldie LC, Nix MK, Hirschi KK. Embryonic vasculogenesis and hematopoietic specification. Organogenesis. 2008;4(4):257–263.

55. Seita J, Weissman IL. Hematopoietic stem cell: self-renewal versus differentiation. Syst Biol Med. 2010;2:640–653.

56. Zonghong S, Meifeng T, Huaquan W, et al. Circulating myeloid dendritic cells are increased in individuals with severe aplastic anemia. Int J Hematol. 2011;156–162.

57. Yu H, Zhao Y, Pan X, Liu C, Fu R. Upregulated expression of profilin1 on dendritic cells in patients with severe aplastic anemia. Front Immunol. 2021;12:1–7.

58. Yankai Xiao, Suwen Zhao, Bo Li. Aplastic anemia is related to alterations in T cell receptor signaling. Stem Cell Investig. 2017;4–85.

59. Frost JN, Wideman SK, Preston AE, et al. Plasma iron controls neutrophil production and function. Sci Adv. 2022;8:eabq5384.

60. Lummertz da Rocha E, Kubaczka C, Sugden WW, et al. CellComm infers cellular crosstalk that drives haematopoietic stem and progenitor cell development. Nat Cell Biol. 2022;24(4):579–589.

61. Vink CS, Calero-Nieto FJ, Wang X, et al. Iterative single-cell analyses define the transcriptome of the first functional hematopoietic stem cells. Cell Rep. 2020;31(6):107627.

62. Mills EW, Wangen J, Green R, Ingolia NT. Dynamic regulation of a ribosome rescue pathway in erythroid cells and platelets. Cell Rep. 2016;17(1):1–10.

63. Kaestner L, Martín-Vasallo P, de La Laguna U, et al. From erythroblasts to mature red blood cells: organelle clearance in mammals. Front Physiol. 2017;8:1076.

64. Zheng Z, Yang S, Gou F, et al. The ATF4-RPS19BP1 axis modulates ribosome biogenesis to promote erythropoiesis. Blood. 2024;144(7):742–756.

65. Xie S, Yan B, Feng J, et al. Altering microtubule stability affects microtubule clearance and nuclear extrusion during erythropoiesis. J Cell Physiol. 2019;19833–19841.

66. Chandrakanthan V, Rorimpandey P, Zanini F, et al. Mesoderm-derived PDGFRA+ cells regulate the emergence of hematopoietic stem cells in the dorsal aorta. Nat Cell Biol. 2022;1211–1225.

67. Méndez-Ferrer S, Michurina T V., Ferraro F, et al. Mesenchymal and haematopoietic stem cells form a unique bone marrow niche. Nature. 2010;466(7308):829–834.

