## Supplementary methods for "Impaired erythroid maturation in murine embryos upon loss of the preeclampsia-associated serine protease prostasin"

Supplementary methods 

FACS analysis
The single cell suspension (as obtained above) was stained with following antibodies: FITC Ter-119 (557915, BD Pharmingen); PE/Cy7 CD71 ab (113812, BioLegend); FITC CD41 ab (561849, BD Pharmigen); PE/Cy7 CD43 ab (121218, BioLegend); BUV737 CD45.2 ab (1313859, BD Horizon); PE CD31 ab (12-0311-82, invitrogen); A700 c-kit ab (56-1172-82, invitrogen); live/dead aqua staining (Invitrogen). Other antibodies used for macrophages, neutrophils, NK cells, T- and B- cells were the following: BV650 CD3e (AB_2738779, BD Horizon); FITC NK-1 (AB_394676, BD Pharmigen); PerCP/Cy5.5 CD11c (117328, Biolegend); BV421 CD11b (101251, Biolegend); PE-texas red F4/80 (AB_10372055, Invitrogen); APC/Cy7 Ly6G (560600, BDPharmigen); BV711 B220 (AB_2738470, BD Horizon). Fluorescent signal was recorded with the DIVA acquisition software using FACS Fortessa machine. The analysis was performed with FlowJo software (Version 10.8.1) for FACS analysis of HSCs and ECs 1, myeloid and lymphoid cells.  

Imaging flow cytometry
Samples were acquired on a 2 camera, 12 channel ImageStreamX imaging flow cytometer (Cytek Biosciences) at low speed and highest magnification (60X). Cells were excited using a 405 nm laser (25mW), a 488 nm laser (150mW), a 561 nm laser (150mW), a 642 nm laser (120mW) and a 785nm Side Scatter (SSC) laser (3 mW). Only events with a bright field area greater than 1 µm² (to exclude cell debris) and non-saturating pixels (Raw max pixel values below 4096) were collected. Data was acquired for at least 10000 events/sample. Experimental samples contained images for bright field (Channels 1 and 9), Ter119-FITC (Channel 2), CD71-PECy7 (Channel 6), Hoechst 33342 (Channel 7), and Side Scatter (Channel 12). Single color controls for each fluorochrome were acquired to generate the compensation matrix that was applied to all the experimental files prior to analysis using IDEAS (Image Data Exploration and Analysis Software) 6.2 software (Cytek Biosciences). For the data analysis of placenta and yolk sac, we first selected two different populations based on Hoechst. Then, Hoechst+ or Hoechst- CD71+cells were selected based on CD71 intensity greater to 104, while an intensity greater than 105 were considered dead cells. Moreover, cell images were analyzed to confirm cell morphology and staining diffusion. Dead cells had high diffusion of Ter119 and CD71 antibodies, suggestive of membrane permeability. For the data analysis of fetal liver samples, we referred to a previous publication 2.
Quantitative reverse transcription PCR (qRT-PCR)
Tissues were flash frozen at -80°C after dissection. For RNA extraction, the tissue was disrupted using metallic balls and TissueLyser at 30 mut/sec for 15 seconds and RNA was isolated using the RNeasy Mini Kit (Qiagen). RNA quality was assessed using Nanodrop spectrometry and 1 µg of RNA was converted into cDNA using the Takara Taq DNA polymerase according to manufacture’s instructions. Gene expression was determined using SYBER Green detection system and the following primers, Prss8: For, 5’- TGACCATTCTGCTCCTTCTC-3’, Rev, 5’- GACACCACCCATTTATTTGAC-3’; Gapdh: For, 5’- CCACCCAGAAGACTGTGGAT-3’, Rev, 5’- CACATTGGGGGTAGGAACAC-3’. Controls using RNA-free water were negative for targets and housekeeping genes. Fold change was calculated using ΔCt method with Gapdh as an internal standard.

Bulk RNA-sequencing and analysis 
RNA was extracted as for qRT-PCR and RNA quality was assessed on a Fragment Analyzer (Agilent Technologies). The RNAs had RQNs between 8.7 and 10. RNA-seq libraries were prepared from 500 ng of total RNA with the Illumina TruSeq Stranded mRNA reagents (Illumina). Library preparation was performed on a Sciclone liquid handling robot (PerkinElmer) with a PerkinElmer-developed automated script. Unique dual indexes were used for barcoding of the libraries. Libraries were quantified by a fluorometric method (QubIT, Life Technologies) and their quality assessed on a Fragment Analyzer (Agilent Technologies). Sequencing was performed on an Illumina NovaSeq 6000. Sequencing data were demultiplexed using the bcl2fastq2 Conversion Software (version 2.20, Illumina). 
For data analysis, FASTQ files were aligned to the mouse genome annotation (GRCm39 release 105, retrieved from Ensembl database) using STAR algorithm (version 2.7.10a) 3. The output count matrices were imported in R (version 4.0.2). AGM and yolk sac data sets were analyzed separately. Analysis of the differentially expressed genes (DEGs) was performed with DESeq2 (version 1.21.1) 4 and p adjusted value of < 0.05 was considered significant. Heatmap representations were made with the ComplexHeatmap package (version 2.4.3) using normalized counts as input. DEGs analysis for the bulk-RNA seq data was performed independently for each genotype comparing time points (E10.5 vs E11.5). The list of DEGs (p adjusted value < 0.05) was divided in upregulated (log2 fold change > 0) and downregulated (log2 fold change < 0) gene sets and subsequently combined in a binary matrix. Uniquely and shared gene sets were displayed as an UpSet plot. 
Functional analysis was performed with the over-representation test function from ClusterProfiler package (version 3.16.1) 5, selecting biological processes as the ontology and using as input specific gene sets derived from the UpSet plots. GO terms were considered significant using a threshold of p adjusted value < 0.05. Heatmap representations for selected GO terms or gene sets were made with the ComplexHeatmap package (version 2.4.3) using normalized counts as input.

Single-cell (sc) RNA-sequencing sample processing
A Chromium Next GEM Chip G (10X genomics) was loaded with the appropriate number of cells, and the sequencing libraries prepared with the Chromium Single Cell 3'reagent kit v3.1 dual index, strictly following the manufacturer’s recommendations. Briefly, an emulsion encapsulating single cells, reverse transcription reagents and cell barcoding oligonucleotides was generated. After the actual reverse transcription step, the emulsion is broken and double stranded cDNA generated and amplified in a bulk reaction (12 PCR cycles). This cDNA was fragmented, a P7 sequencing adaptor ligated, and a 3’ gene expression library generated by PCR amplification (13 cycles). Libraries were quantified by a fluorometric method, and their quality assessed on a Fragment Analyzer (Agilent Technologies). Sequencing was performed on Illumina NovaSeq 6000 v1.5 flow cells for 28-10-10-90 cycles (read1 - index i7 - index i5 - read2) with 1% PhiX spike in. 

scRNA-sequencing data analysis and bioinformatics
Fastq files were processed with CellRanger count with the default settings (cellranger-7.0.1) on the mouse genome reference refdata-gex-mm10-2020-A. Count matrices were imported into R (version 4.0.2) using the Seurat package (version 4.0.1) 6. Cell filtering was performed by applying sample specific thresholds after exploratory analysis. More specifically, cells with more than 1000 (AGM E10.5 dataset) or 1500 (AGM E11.5 dataset) and less than 7000 features, and mitochondrial percentage lower than 10% were kept for further analysis. Both Seurat objects were then combined using RPCA integration. Cell type annotation was performed with singleR package (version 1.2.4) 7 using as reference a subset of the scRNA seq dataset derived from mouse gastrulation and early organogenesis. 8 More specifically we used the set derived from the E10.25 stage, the latest stage present in the dataset and the closest to the time points included in our experiment. Additionally, the automatic cell annotations were manually adjusted and refined based on reported gene markers 9. DEGs analyses were performed using Seurat (version 4.2.0)10 built-in functions (FindConservedMarkers, FindAllMarkers). Moreover, differentially expressed genes with a p adjusted value <0.05 were considered significant and used as input for other downstream analyses as required. Functional analysis of gene sets was performed with the over-representation test function from ClusterProfiler (version 3.16.1) selecting biological processes as the ontology. 
For entropy estimation each tissue data set was divided by time point, while keeping both genotypes in each subset. Single cell entropies were computed using the SLICE algorithm (version 0.99.0)11 and the results were visualized with ggplot2 (version 3.3.3).
Analysis of the Diamond-Blackfan anemia genes was performed using as reference a list of human genes retrieved from the French National Diagnosis and Care Protocol (PNDS).12 The human gene names were translated to the mouse orthologs with the biomaRT package (version 2.44.4) and then intersected with the sets of DEGs of interest. 
Heatmap representations for selected GO terms or gene sets were made with the ComplexHeatmap package (version 2.4.3) using as input normalized counts.

Supplementary References 


1.	Zhou F, Li X, Wang W, et al. Tracing haematopoietic stem cell formation at single-cell resolution Isolation of single T1 pre-HSC by CD201. Nature. 2016;533:487–492. 
2.	McGrath KE, Catherman SC, Palis J. Delineating stages of erythropoiesis using imaging flow cytometry. Methods. 2017;112:68–74. 
3.	Dobin A, Davis CA, Schlesinger F, et al. Sequence analysis STAR: ultrafast universal RNA-seq aligner. BIOINFORMATICS. 2013;29(1):15–21. 
4.	Love MI, Huber W, Anders S. Moderated estimation of fold change and dispersion for RNA-seq data with DESeq2. Genome Biol. 2014;15(12):. 
5.	Yu G, Wang LG, Han Y, He QY. ClusterProfiler: An R package for comparing biological themes among gene clusters. OMICS. 2012;16(5):284–287. 
6.	Satija R, Farrell JA, Gennert D, Schier AF, Regev A. Spatial reconstruction of single-cell gene expression data. Nat Biotechnol. 2015;33(5):495–502. 
7.	Aran D, Looney AP, Liu L, et al. Reference-based analysis of lung single-cell sequencing reveals a transitional profibrotic macrophage. Nat Immunol. 2019;20:163–172. 
8.	Pijuan-Sala B, Griffiths JA, Guibentif C, et al. A single-cell molecular map of mouse gastrulation and early organogenesis. Nature. 2019;566:490–495. 
9.	Lummertz da Rocha E, Kubaczka C, Sugden WW, et al. CellComm infers cellular crosstalk that drives haematopoietic stem and progenitor cell development. Nat Cell Biol. 2022;24(4):579–589. 
10.	Butler A, Hoffman P, Smibert P, Papalexi E, Satija R. Integrating single-cell transcriptomic data across different conditions, technologies, and species. Nat Biotechnol. 2018;36(5):411–420. 
11.	Guo M, Bao EL, Wagner M, Whitsett JA, Xu Y. SLICE: determining cell differentiation and lineage based on single cell entropy. Nucleic Acids Res. 2017;45(7):54. 
12.	Member of the MaRIH rare diseases health network. French National Diagnosis and Care Protocol (PNDS): acquired and inherited aplastic anemia. 2023. 
 
